# Chondroitin Sulfate Elicits Systemic Pathogenesis In Mice By Interfering With Gut Microbiota Homeostasis

**DOI:** 10.1101/142588

**Authors:** Tao Liao, Yan-Ping Chen, Shui-Qing Huang, Li-Li Tan, Chang-Qing Li, Xin-An Huang, Qin Xu, Qi Wang, Qing-Ping Zeng

## Abstract

Whether chondroitin sulfate (CS), a common ingredient naturally occurring in livestock and poultry products, improves osteoarthritis remains debating. Here, we show for the first time that CS induces steatogenesis, atherogenesis, and dementia-like pathogenesis in mice. Gut microbiome analysis revealed the sulfatase-secreting bacteria *Rikenella* and the sulfate-reducing bacteria *Desulfovibrio* are enriched. Surprisingly, berberine use boosts CS-induced multi-loci inflammatory manifestations by further increasing the abundance of *Rikenella* and *Desulfovibrio*, whereas cephalosporin reinforces the colon mucus barrier via flourishing *Akkermansia muciniphila* and upregulating mucin expression. Mechanistically, berberine aggravates mucus lining injury by prompting mucin degradation, endotoxin leakage, neutralizing antibody induction, pro-inflammatory cytokine burst, lactic acid accumulation and energy currency depletion in multiple organs and tissues. Taken together, CS evokes the early-phase pathogenesis toward steatohepatitis, atherosclerosis, and dementia upon augmenting gut opportunistic infection, and a sustained antibiotic monotherapy does not deprive the risk of CS-driven systemic inflammatory disorders.

## INTRODUCTION

Chondroitin sulfate (CS), a common biochemical ingredient in livestock and poultry products, is prescribed as a drug for treatment of osteoarthritis (OA) in 22 countries, including the European Union, based on the efficacy and safety data available from clinical trials (Vergés et al., 2004). Nevertheless, CS is still regulated in the United States as a dietary supplement. In 2004, the Food and Drug Administration (FDA) denied a petition that requests CS be labeled as reducing the risk of OA and OA-related joint pain, tenderness, and swelling (FDA, 2004). A systematic review summarizing 20 clinical trials concluded the symptomatic benefit of CS from the large-scale and methodologically sound trials is minimal or nonexistent, so CS use in the routine clinical practice should be discouraged (Reichenbach et al., 2007). However, a contrast conclusion was drawn that compelling evidence supports CS interferes with OA progression (Clegg et al., 2006; Reginster et al., 2007).

An updated Cochrane review on the clinical trials involving CS indicated most are of low quality, but there is some evidence of short-term improvement in pain and few side effects; it does not appear to improve or maintain the health of affected joints (Singh et al., 2015). it was warned that CS should not be used to treat patients with symptomatic knee OA because of failure in relief of the condition (Jevseva et al., 2013). It was also demonstrated that a combined therapy of CS with glucosamine sulfate or glucosamine hydrochloride does not improve joint damage in a rabbit model of knee OA (Roman-Blas et al., 2017a), and CS plus glucosamine sulfate shows no superiority over placebo in a randomized, double-blind and placebo-controlled clinical trial in patients with knee OA (Roman-Blas et al., 2017b). In contrast, an anti-inflammatory effect of CS was filed in amelioration of some inflammatory diseases, at least including inflammatory bowel disease (IBD) (Segarra et al., 2016; Holleran et al., 2017).

While CS is helpful to OA remains controversial, the inconsistent results regarding the “beneficial” role of CS in OA might be argued not to be attributed to the “bad” design of experiments. Actually, there are challenges against contamination of CS with heparin, which may cause some adverse clinical events, such as allergy and hypertension (Guerrini et al., 2008; Kishimoto et al., 2008). However, no reports was noted to concern a potential effect of the individual gut microbiota on the differential pharmaceutical outcome of CS. Although evidence on CS-driven inflammatory pathogenesis by interfering with the gut microbiota is still lacking, an implication of the gut microbiota disturbance has been addressed in atherosclerosis (Spence, 2016; Wilson et al., 2016), non-alcoholic fatty liver disease (NAFLD) (Mokhtari et al., 2017), non-alcoholic steatohepatitis (NASH) (Brandl & Schnabl, 2017), and cardiovascular diseases (CVD) (Lippi et al., 2017; Garcia-Rios et al., 2017).

As a common ingredient in livestock and poultry products, CS was recognized to increase the abundance of *Bacteroides thetaiotaomicron*, a species of sulfatase-secreting bacteria (SSB) that degrades the glycoprotein-rich dietary sulfate and colon mucus, which supply the sulfate to *Desulfovibrio piger*, a sulfate-reducing bacteria (SRB) (Rey et al., 2013), arising a possibility of CS-driven coordinated overgrowth of SSB and SRB as a food chain. Indeed, many species of CS-degrading bacteria, including *B. thetaiotaomicron* J1 and 82, *B. ovitus* E3 and *Clostridium hathewayi* R4, were isolated from healthy subjects (Shang et al., 2016). Alternatively, heme as a common component in the red meat was found to increase the abundance of *Akkermansia muciniphila*, a mucin-degrading and mucus-damaging commensal gut microbiota, which can facilitate the aberrant colon epithelial proliferation by exploiting the mucin-containing carbohydrates and amino acids as substrate for growth, and also by employing the mucin-derived sulfate as an electron acceptor to give rise to sulfide and sulfur dioxide (Ijssennager et al., 2015).

Considering CS as a sulfate is similar with mucin, a typical protein constituent of the mucus layer overlying the gut epithelium (Johansson et al., 2013), we suggest herein a hypothesis of “organ-specific anti - or pro-inflammation” followed by CS-induced SSB overgrowth, mucin degradation and endotoxin leakage. In a more detail, CS promotes SSB overgrowth, and then SSB degrade the O-glycon-containing mucin, hence opening the mucin-crossing mucus barrier, and leaking the bacterial endotoxin lipopolysaccharide (LPS) into the blood. The circulated LPS would elicit production of neutralizing antibody and secretion of pro-inflammatory cytokines to scavenge LPS, during which organs without LPS might show an anti-inflammatory feature, whereas organs with LPS might exhibit a pro-proinflammatory character. Of course, CS either ameliorates or aggravates OA might be also dependent on the individual gut microbiota status.

We fed mice with 25 mg of CS daily to induce gut opportunistic infection, and also used 100 mg/ml berberine (BER) or 100 mg/ml cephalosporin (CEF) to control gut opportunistic infection. BER is considered antibiotic (Li & Zuo, 2010) albeit with a low bioavailability (Liu et al., 2016), so it should directly and non-selectively kill gut bacteria. Upon fed CS combined with or without BER or CEF, we disclosed for the first time that CS not only flourishes SSB and interferes with gut microbiota homeostasis, but also induces the multiple inflammatory pathogenesis, such as steatogenesis, atherogenesis, and dementia-like amyloidosis and cognitive deficits. Surprisingly, BER use for one month was noticed to aggravate the multiple pathogenic alterations by increasing the richness of SSB. It could be anticipated that the astonishing achievements regarding CS - and BER/CEF-exerted dual effects of anti - and pro-inflammation should pave a path toward a thorough elucidation of the rationale underlying gut opportunistic infection-originated inflammatory disorders and metabolic syndromes.

## RESULTS

### CS Alters Gut Microbiota Communities And BER/CEF Interferes With Gut Microbiota Homeostasis

To assess the effect of animal-based diets on the gut microbiota homeostasis, we fed mice with CS or lard (LD) and performed the 16S VX region-based gut microbiome analysis (Additional file 1). As results, all tested mice show two major gut microbiota phyla among Top 10 abundant phyla: *Firmicutes* and *Bacteroidetes*. As illustrated in Figure 1A, CS nourishes more *Firmicutes* and less *Bacteroidetes*, whereas LD flourishes more *Bacteroidetes* and less *Firmicutes*. By comparison with Top 35 abundant genera, CS mice exhibit 12 abundant genera (red color) within *Firmicutes* and two abundant genera within *Proteobacteria*, whereas LD mice possess two abundant genera within *Firmicutes* and one abundant genus within *Actinobacteria* (Figure 1B). As indicated, while *Bacteroides* are bile tolerant bacteria that adapt to high-fat diets, *Firmicutes* are polysaccharide degradable bacteria that adapt to high-fiber diets (David et al. 2014).

**Figure 1.**
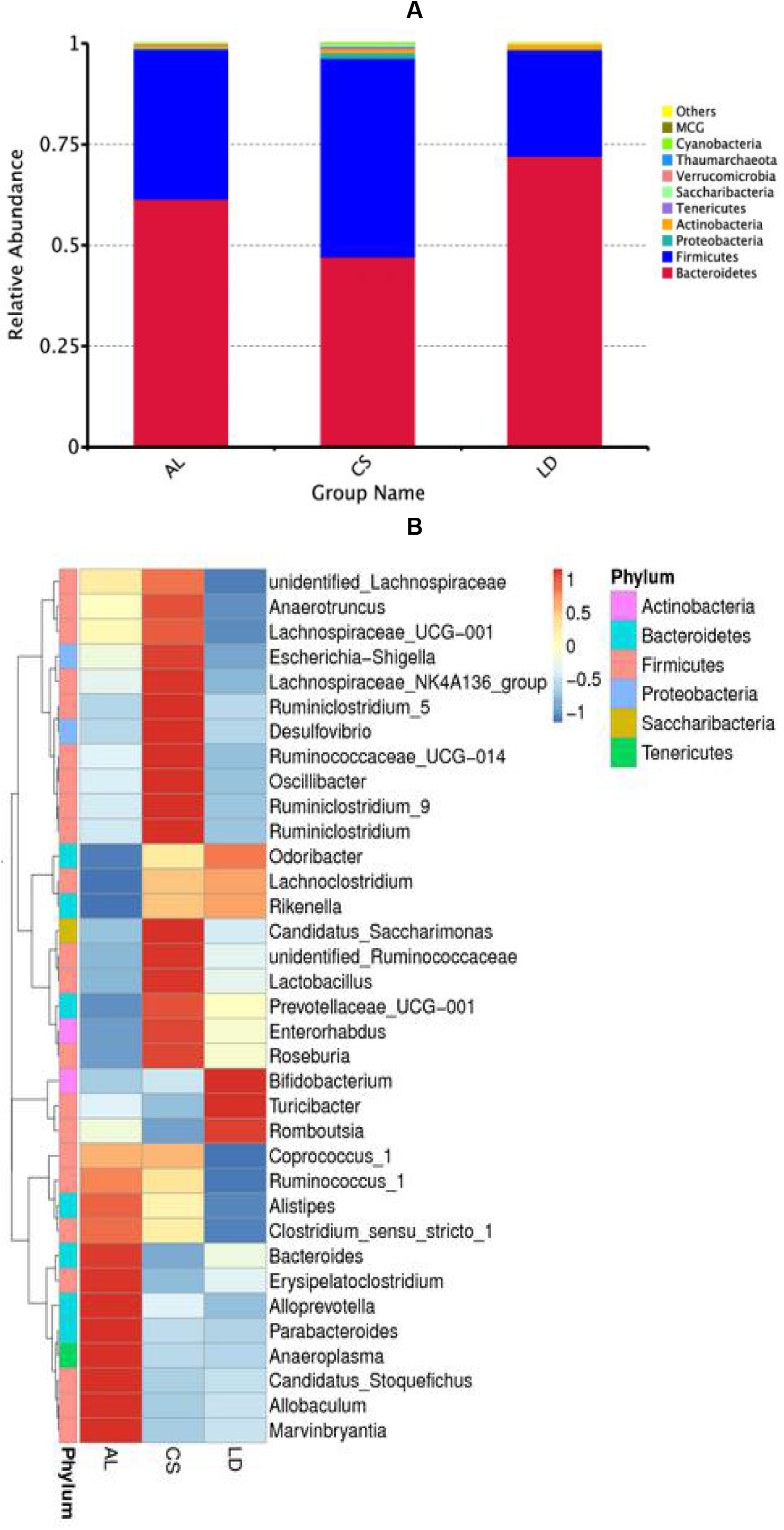
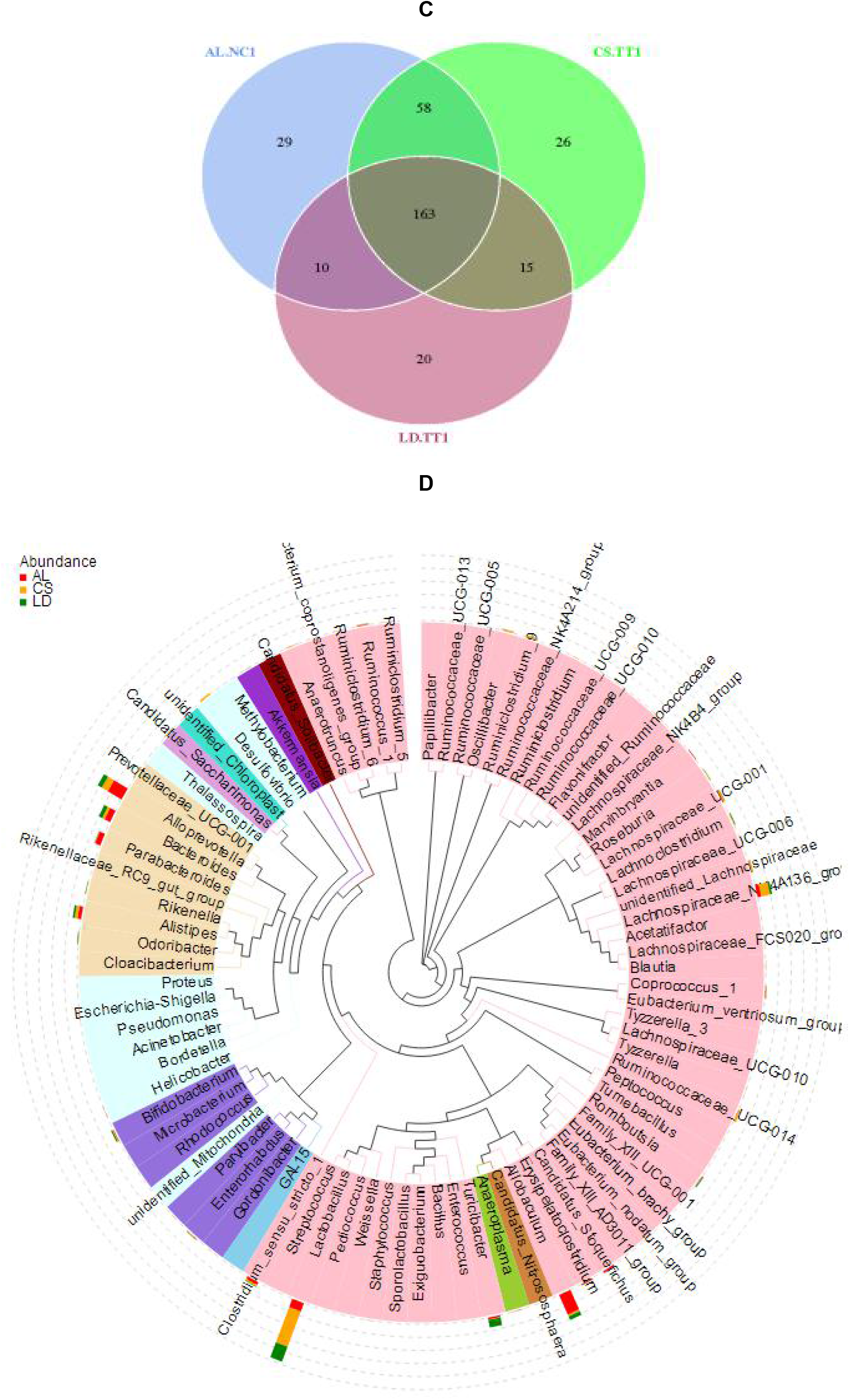
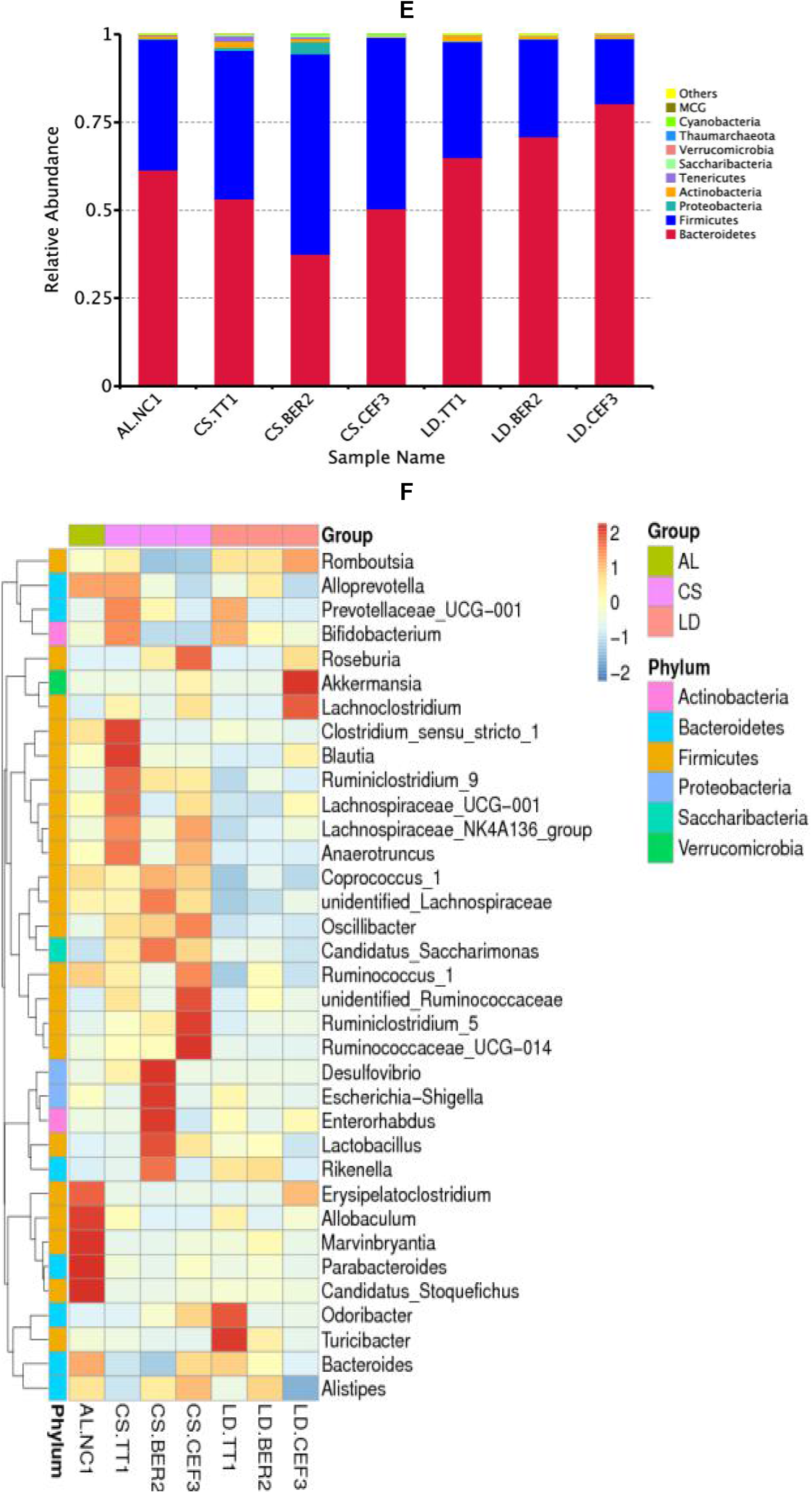
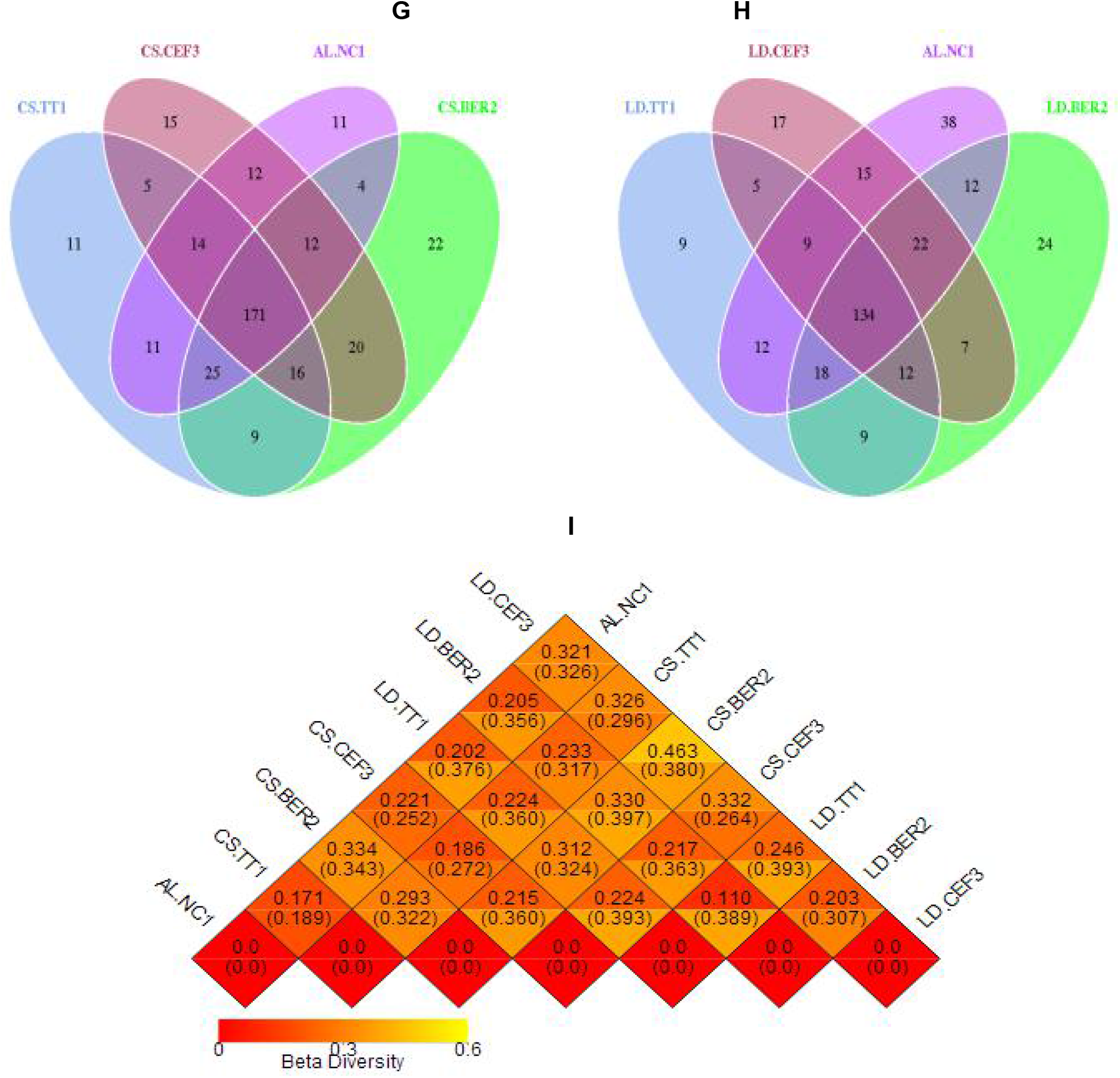
The gut microbiota profiles in AL, CS, LD, BER, and CEF mice. (A) The relative abundance of Top 10 genera in AL, CS, and LD mice. (B) Top 35 abundant genera in AL, CS, and LD mice. (C) Top 100 OTUs in AL, CS, and LD mice. (D) Top 100 abundant genera in AL, CS, and LD mice. (E) The relative abundance of Top 10 genera in AL.NS1, CS.TT1, CS.BER2, CS.CEF3, LD.TT1, LD.BER2, and LD.CEF3. (F) Top 35 abundant genera in AL.NS1, CS.TT1, CS.BER2, CS.CEF3, LD.TT1, LD.BER2, and LD.CEF3. (G)Top 100 OTUs in AL.NS1, CS.TT1, CS.BER2, and CS.CEF3. (H) Top 100 OTUs in AL.NS1, LD.TT1, LD.BER2, and LD.CEF3. (I) β-diversity in AL.NS1, CS.TT1, CS.BER2, CS.CEF3, LD.TT1, LD.BER2, and LD.CEF3.

It is obviously that CS also increases an abundance of the harmful *Escherichia-shigella*, whereas LD allows an overgrowth of the beneficial *Bicidobacterium*. Additionally, Top 100 operational taxonomic units (OTUs) are more similar between AL.NC1 and CS.TT1 than other comparisons because they share 58 common OTUs (Figure 1C), suggesting LD causes more gut microbiome changes than CS. According to the evolutionary tree constructed from Top 100 abundant genera, it was distinguishable that *Lactobacillus* is the most abundant genus in both CS and LD mice, and *Allobacterium* and *Alloprevotella* are the most abundant genera in AL mice (Figure 1D).

To evaluate the effects of antibacterial agents on gut microbiota changes, we treated CS/LD mice with BER or CEF and monitored gut microbiome changes. As compared with CS.TT1, BER significantly increases the abundance of *Firmicutes* in CS.BER2, whereas CEF sightly increases the abundance of *Firmicutes* in CS.CEF3 (Figure 1E). In contrast, while BER slightly increases the richness of *Bacteroidetes*, CEF significantly increases the richness of *Bacteroidetes* Surprisingly, BER enriches *Desulfovibrio*, as seen from CS.BER2, and CEF flourishes *A. muciniphila*, as seen from LD.CEF3 (Figure 1F). It was observed that 171 common OTUs are shared among AL.NC1, CS.TT1, CS.BER2, and CS.CEF3 (Figure 1G). On the other hand, it was also noticed that there are 134 common OTUs among AL.NC1, LD.TT1, LD.BER2, and LD.CEF3 (Figure 1H). As considering the β-diversity that represents the proportion between regional and local species diversity, it was seen that AL.NC1 has the least difference from CS.TT1, but the most difference from LD.CEF3 (Figure 1I).

Our determination revealed CS enriches 53% of *Bacteroidetes* and 42% of *Firmicutes*. In similar, LD enriches 65% of *Bacteroidetes* and 33% of *Firmicutes*. These results addressed livestock and poultry-sourced CS and LD allow the overgrowth of *Bacteroidetes*. Other authors also indicated an animal-based high fat/high protein food intake can increase the abundance of bile tolerant *Bacteroides, Alistips* and *Bilophila*,whereas a plant-based high fiber food intake can increase the richness of polysaccharide degradable *Firmicutes* and *Prevotella* (Wu et al., 2011; David et al., 2014; Kovatcheva-Datchary et al., 2015).

### CS/LD And BER/CEF Enrich Different Bacterial Genera/Species Belonging To SRB And SSB

To investigate the difference and extent of CS/LD and BER/CEF in enriching SRB and SSB, we particularly analyzed the proportion of *Desulfovibrio, Rikenella*, and *A. muciniphila* in the Kingdom of *Bacteria* (Table 1). It was noticeable that CS increases the abundance of *Desulfovibrio* from 0.006% (2% in the Phylum of *Proteobacteria*) in AL.NC1 (Figure 2A) to 0.70% (92% in *Proteobacteria*) in CS.TT1 (Figure 2B), accounting for more than 100-fold increases. While BER further increases its abundance to 3.00% (74% in *Proteobacteria*) in CS.BER2 (Figure 2C), CEF decreases its abundance to 0.003% (40% in *Proteobacteria*) in CS.CEF3 (Figure 2D). It was also seen from Table 1 that CS increases the abundance of *Rikenella* from 0.001% in AN.NC1 to 0.05% in CS.TT1, estimating up to 50-fold increases. BER further increases its abundance to 0.70% in CS.BER2, whereas CEF decreases its abundance to 0 in CS.CEF.

**Figure 2.**
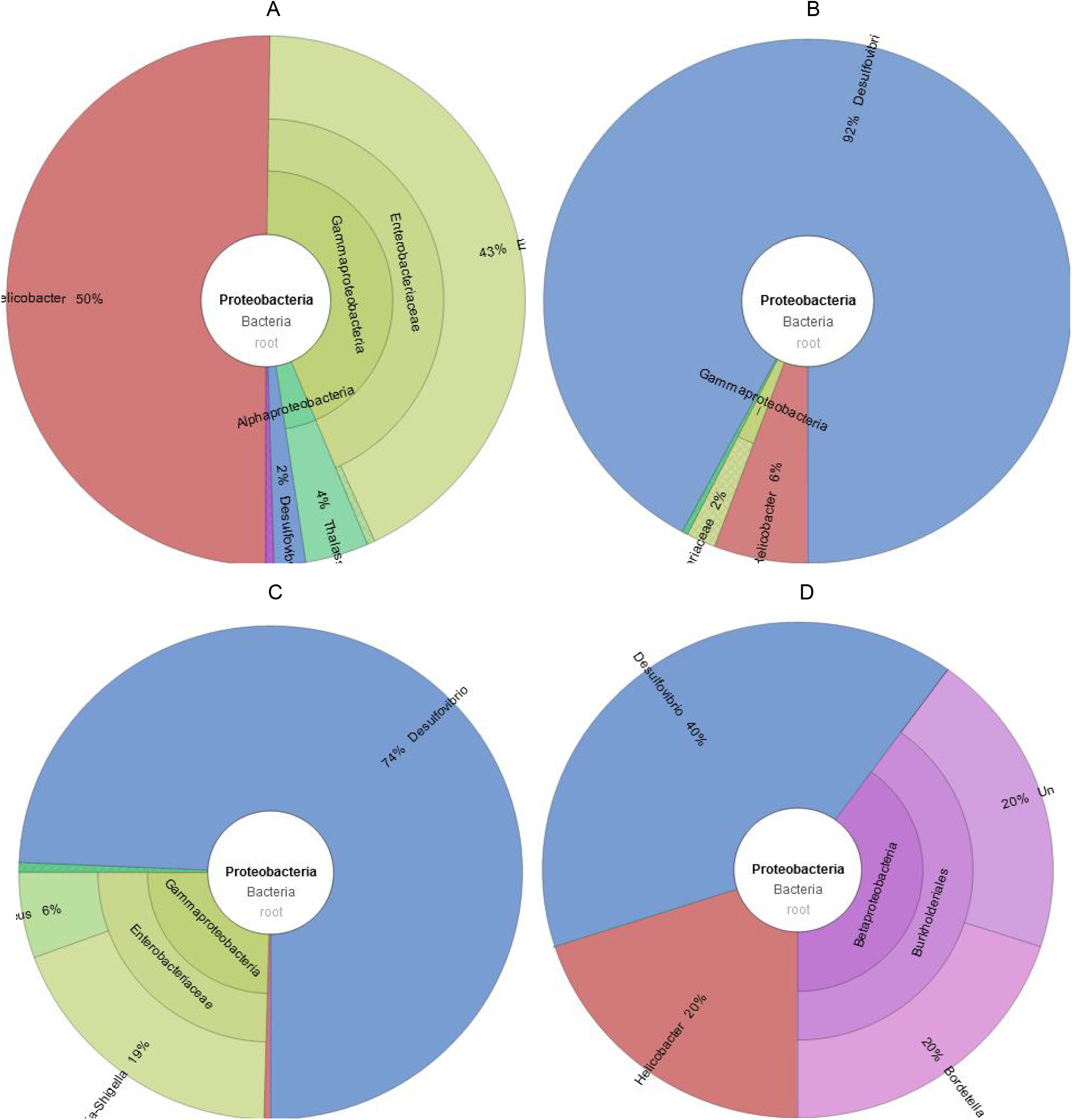
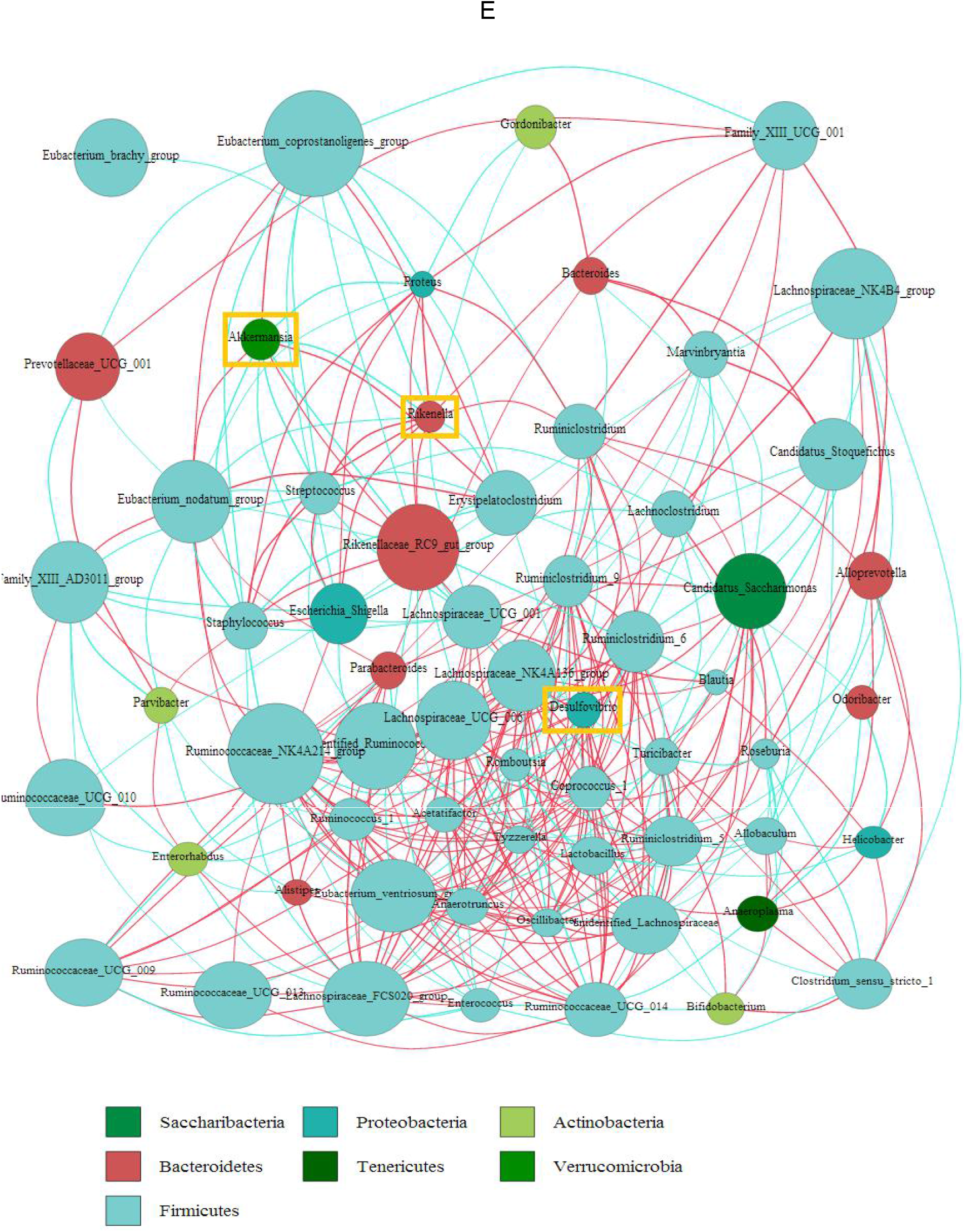
The abundance of *Desulfovibrio* in *Proteobacteria* and interactions among *Desulfovibrio, Rikenella*, and *Akkermansia*. (A) The percentage of *Desulfovibrio* in AL.NC1. (B) The percentage of *Desulfovibrio* in CS.TT 1. (C) The percentage of *Desulfovibrio* in CS.BER2. (D) The percentage of *Desulfovibrio*in CS.CEF3. (E) An interactive network among *Desulfovibrio, Rikenella*, and *Akkermansia* (yellow boxes), in which the size of a dot represents an average relative abundance in a specific genus, and the line between every dot indicates a positive (red) or negative (blue) correlation.

**Table 1.** The proportion of selective bacterial phyla and genera in AL, CS, LD, BER, and CEF mice.

| Phyla/Genera | AL.NC1 | CS.TT1 | CS.BER2 | CS.CEF3 | LD.TT1 | LD.BER2 | LD.CEF3 |
| --- | --- | --- | --- | --- | --- | --- | --- |
| Phylum: *Proteobacteria* | 0.30 | 0.80 | 3.00 | 0.007 | 0.20 | 0.09 | 0.04 |
| Genus: *Desulfovibrio* | 0.006 | 0.70 | 3.00 | 0.003 | 0 | 0 | 0 |
| Genus: *Escherichia-shigella* | 0.10 | 0.009 | 0.70 | 0 | 0.20 | 0.03 | 0 |
| Phylum: *Verrucomicrobia* | NT | NT | NT | NT | NT | NT | NT |
| Species: *Akkermansia muciniphila* | 0.01 | 0.01 | 0.00 | 0.09 | 0.00 | 0.00 | 0.30 |
| Phylum: *Bacteroidetes* | 61.00 | 53.00 | 38.00 | 50.00 | 65.00 | 71.00 | 80.00 |
| Genus: *Rikenella* | 0.001 | 0.05 | 0.70 | 0 | 0.40 | 0.40 | 0 |
| Phylum: *Firmicutes* | 37.00 | 42.00 | 57.00 | 49.00 | 33.00 | 28.00 | 18.00 |
| Genus: *Lactobacillus* | 6.00 | 8.00 | 43.00 | 24.00 | 13.00 | 16.00 | 3.00 |
| Phylum: *Actinobacteria* | 0.80 | 2.00 | 0.90 | 0.10 | 2.00 | 0.90 | 0.80 |
| Genus: *Bifidobacterium* | 0.50 | 2.00 | 0.00 | 0.003 | 1.00 | 0.70 | 0.40 |

Note: The proportion is represented by the percentage (%). NT: not tested.

In the LD groups, it was observed that LD, BER or CEF does not support the growth of *Desulfovibrio*, which is undetectable in LD.TT1, LD.BER2 and LD.CEF3 (Table 1). As to *Rikenella*, its abundance is increased from 0.001% in AL.NC1 to 0.40% in LD.TT1, accounting for an increase of 400 folds, suggesting LD prompts the growth of *Desulfovibrio*. Regarding the effect of antibiotics, it was measured that LD.BER2 harbors 0.40% of *Desulfovibrio*, whereas LD.CEF3 decreases its abundance to 0, indicating CEF rather than BER can eradicate *Desulfovibrio* (Table 1). A similar trend was also observed in the CS group, where BER increases the abundance of *Desulfovibrio*, while CEF decreases its abundance.

As a species belonging to mucin-degrading bacteria (MDB), *A. muciniphila* was not observed to be induced by CS because its abundance in CS.TT1 is the same as that in AL.NC1 (0.01%). Surprisingly, CEF increases its abundance from 0.01% in CS.TT1 to 0.09% in CS.CEF3 as well as from 0 in LD.TT1 to 0.3% in LD.CEF. In contrast, BER can eradicate *A. muciniphila* (Table 1). These results highlighted *A. muciniphila* might only “eat” mucin but not “eat” CS, which means *A. muciniphila* is unlikely a CS-induced species of SSB. However, *A. muciniphila* might be survived and even flourished by antibiotics, such as CEF, via killing other bacteria.

Besides, BER also increases an abundance of the pathogen *Escherichia-shigella* from 0.10% in AL.NC1 to 0.70% in CS.BER2 or 0.20% in LD.TT1, but decreases its abundance to 0.001% in CS.TT1, 0.03% in LD.BER2 or0 in LD.CEF3. Intriguingly, BER extremely increases the abundance of *Lactobacillus*, one genus of probiotic bacteria, but eradicate *Bifidobacterium*, another genus of probiotic bacteria (Table 1).

Although an interaction of the gut microbiota is complex, it was still aware that *Desulfovibrio* and *Rikenella* are positively or dependently correlated, whereas *Rikenella*and *Akkermansia* are negatively or independently correlated (Figure 2E). It should be understandable because *Rikenella* as SSB provide sulfate to *Desulfovibrio* as SRB, but both *Rikenella* and *Akkermansia* are SSB.

CS exhibits a growth inhibitory effect on *Bacteroidetes* by decreasing its abundance from 61% to 53%, including decreasing the abundance of *Bacteroides* from 4% to 1%. While CEF eradicates the pathogen *Escherichia-shigella* and also decreases the abundance of the probiotic *Bifidobacterium*, BER cannot eradicate *Escherichia-shigella*, but can kill *Bifidobacterium*. It was evident that *Desulfovibrio* use sulfate to generate hydrogen sulfide, which can be convert to thiosulfate and further oxidized to tetrathionate, thereby promoting the overgrowth of γ-proteobacteria, including *Escherichia-shigella* (Rajilić-Stojanović et al.2015) Briefly, CS enriches *Desulfovibrio* and *Rikenella*, by which a metabolic chain that links O-glycan sulfate induction to sulfatase secretion, sulfate release and sulfate reduction is formed. BER potentiates such an association by further enriching *Desulfovibrio* to 3% and *Rikenella* to 2%. Additionally, *A. muciniphila* is enriched by CEF but not by CS and LD.

It was previously confirmed that SRB are colonized in the human gut accounting for a proportion of about 50% among the commensal microbes (Ijssennager et al., 2015), including 66% of *Desulfovibrio* and 16% of *Desulfobulbus* (Leschelle et al., 2005). Although an almost equal proportion of SRB might be anticipated in this study, we only identified *Desulfovibrio* in AL.TT1 and CS.TT1 for 0.01% and 0.70%, suggesting a potential existence of SRB in the unidentified SSB (Rey et al., 2013). SRB exploit sulfate as a terminal electron acceptor in their respiratory chains to oxidize the organic compounds and hydrogen into hydrogen sulfide, which can directly induce free radicals, prompt DNA damage, and increase the risk of colon cancer (Attene-Ramos et al., 2007). For example, *A. muciniphila* as SSB and SRB can reduce sulfate to sulfide (Ijssennager et al., 2015).

### Colon Lesions Are Dependent Upon *Desulfovibrio* And *Rikenella* But Colon Repairs Are Enhanced By *A. muciniphila*

To assess a causal linkage of SSB/SRB to mucus layer integrity, we detected the secretary leukocyte protease inhibitor (SLPI), a hallmark of colon lesions (Clauss et al., 2002), in the descending colon lining of all tested mice. As compared with AL.NC1 (Figure 3A1), the color that correlates with the SLPI level was discernible to be dark brown in CS.TT1 (Figure 3A2) and darker brown in CS.BER (Figure 3A3), but lighter brown in CS.CEF (Figure 3A4). Among the LD group, their colon linings generally show a relatively lower SLPI level than the CS group, in which LD.CEF3 shows the lowest SLPI level (Figure 3A5-3A7).

**Figure 3.**
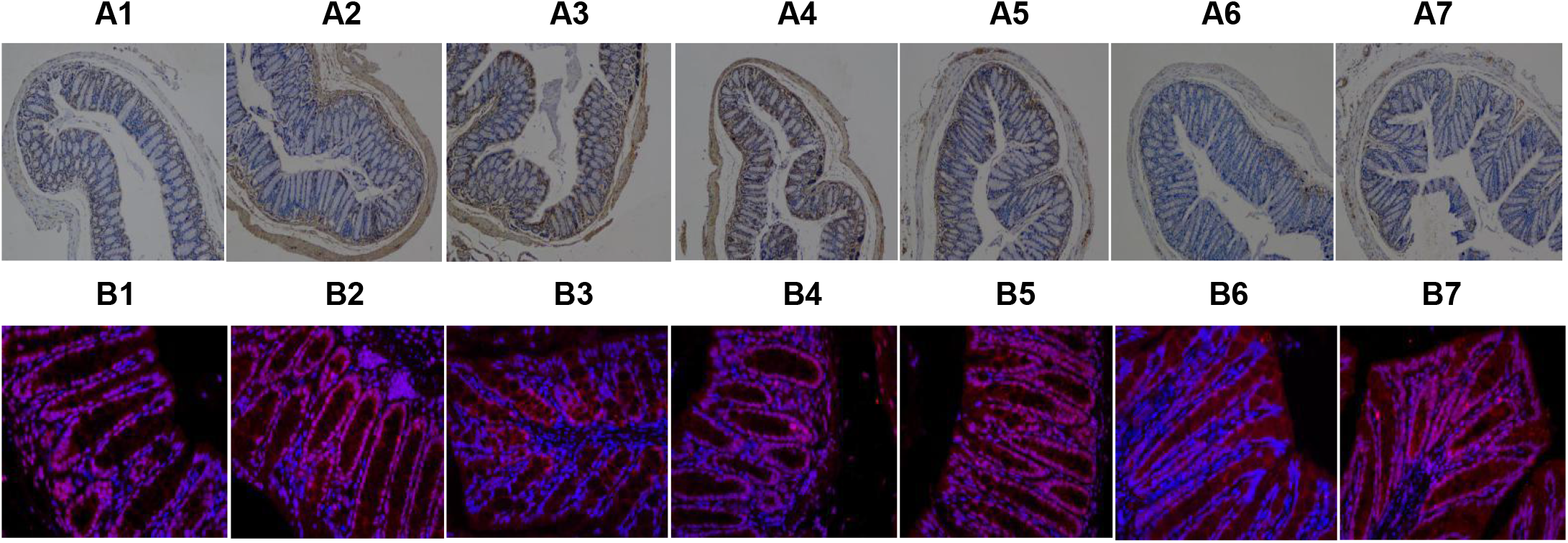
Expression of SLPI and MUC1 in the colon linings of AL, CS, LD, BER, and CEF mice. (A) SLPI staining (× 200). (B) MUC1 fluorescence (× 400). A1-A7 and B1-B7 represent AL.NC1, CS.TT1, CS.BER2, CS.CEF3, LD.TT1, LD.BER2, and LD.CEF3, respectively. SLPI was stained into the brown color around the colon linings. The red fluorescence represents MUC1, and the blue fluorescence represents DNA stained by DPCI.

Interestingly, the overexpression of SLPI is likely correlated with the abundance of *Desulfovibrio* and *Rikenella* (see Table 1). For example, CS.TT1 and CS.BER2 with abundant *Desulfovibrio* and *Rikenella* show the higher SLPI levels than AL.NC1. In contrast, CS.CEF3 with trace-number *Desulfovibrio* but without *Rikenella* shows the lower SLPI levels than CS.TT1 and CS.BER2. In similar, LD.TT1 and LD.BER2 with trace-number *Rikenella* but without *Desulfovibrio* also exhibit the lower SLPI levels. In particular, LD.CEF3 without *Desulfovibrio* and *Rikenella* show the lowest SLPI levels.

As compared with AL.NC1 (Figure 3B1), it was clearly that the pink dots representing the merged blue fluorescence of DNA and red fluorescence of mucin 1 (MUC1) are much less in the colon mucus linings of CS.BER2 than those of CS.TT1 and CS.CEF3, implying a dramatic downregulation of MUC1 in CS.BER2. At the same time, the colon mucus linings of CS.BER2 also exhibit many mesh-like structures with huge holes (Figure 3B2-3B4), also implying an extreme downregulation of MUC1 and other subtypes of mucins in CS.BER2. Interestingly, CS.BER2 does not harbor *A. muciniphila*, whereas AL.NC1, CS.TT1 and CS.CEF3 carry *A. muciniphila* (see Table 1), suggesting a possible correlation of MUC1 overexpression and colon repairs with *A. muciniphila* presence.

In contrast, it was unambiguously distinguished that LD.CEF3 possesses much more pink dots and much less mesh-like structures with huge holes than LD.TT1 and LD.BER2 (Figure 3B5-3B7), indicating an overexpression of MUC1 in LD.CEF3. Coincidentally, LD.CEF3 harbors abundant *A. muciniphila*, whereas LD.TT1 and LD.BER2 does not carry *A. muciniphila* (see Table 1).

The gut bacteria MDB/SSB are classified into two categories: one is mucin specialists that only grow on mucin O-glycans as a sole polysaccharide source (such as *A. muciniphila* and *Barnesiella intestinihominis*), and another is mucin generalists that grow on several other polysaccharides (such as *B. thetaiotaomicron* and *Bacteroides* caccae) (Desai et al., 2016). Using the gut microbiota metagenomic analysis, we identified *A. muciniphila* and *Rikenella*. Nevertheless, it should be predicted that some other MDB/SSB might be included in the unidentified *Bacteroidetes*, which exist in LD mice for 65% and in CS mice up to 53%.

### Endotoxin Leakage Leads To Prompted LPS Circulation and Accelerated Anti-LPS Induction

Because of colon mucus openings, LPS derived from Gram negative bacteria can be readily leaked into the blood stream and induce anti-LPS induction. To evaluate the interactive changes of LPS and anti-LPS, we measured at first their peripheral (serum and muscular) levels. As shown in Figure 4A-4B, it was unexpectedly observed that the serum and muscular LPS levels are almost declined among all treated mice, even lower than AL mice, suggesting CS/LD might elevate LPS levels at first, then induce anti-LPS to neutralize LPS, and eventually decline LPS levels. Subsequently, we determined the visceral (adipose, hepatic, cardiac and cerebral) LPS levels (Figure 4C-4F). It was seen that the multiple organ LPS levels in treated mice are also nearly equal to or lower than those in AL mice, also suggesting a trend of an elevated LPS level at first and then a declined LPS level in CS/LD mice.

**Figure 4.**
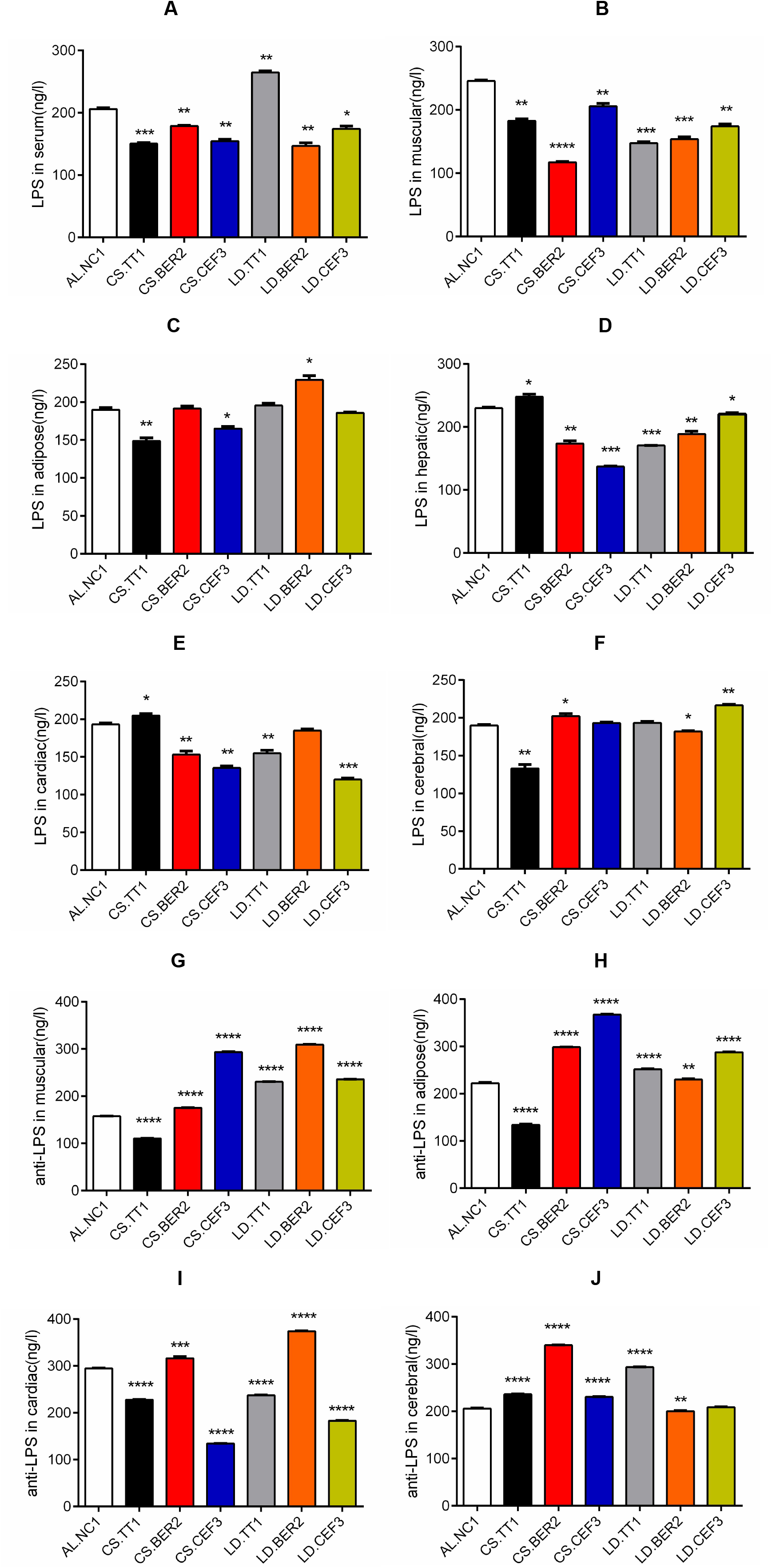
The serum and muscular levels of LPS, anti-LPS and TNF-α in AL, CS, LD, BER, and CEF mice. (A) LPS levels in the serum. (B) LPS levels in the muscular tissue. (C) LPS levels in the adipose tissue. (D) LPS levels in the hepatic tissue. (E) LPS levels in the cardiac tissue. (F) LPS levels in the cerebral tissue. (G) Anti-LPS levels in the muscular tissue. (H) Anti-LPS levels in the adipose tissue. (I) Anti-LPS levels in the cardiac tissue. (J) Anti-LPS levels in the cerebral tissue.

As evidence supporting the above prediction that LPS induces anti-LPS at first, anti-LPS secondly neutralizes LPS, and eventually declines the LPS levels, it was obviously that the multiple organ anti-LPS levels in all treated mice are generally equal to or higher than those in AL mice. Furthermore, it was clearly that antibiotic use, of course, affects the dynamic interaction between LPS and anti-LPS as antibiotics might decline or elevate the LPS levels. The muscular, adipose, cardiac, and cerebral anti-LPS levels (Figure 4G-4J) in CS.BER2 are higher than AL.NC1. These results indicated an active immunostimulatory response is triggered in CS.BER2 to give rise to an adverse inflammatory outcome.

### Systemic Inflammation Is Followed By Upregulated Mitochondrial Biomarkers

To monitor whether LPS systemically induces the pro-inflammatory responses, we measured the peripheral levels of tumor necrosis factor α (TNF-α), a typical and major pro-inflammatory cytokine. Consequently, TNF-α maintains a relatively low level in the serum among almost all groups of mice except for LD.TT1, but a high level in the muscular tissue in CS.TT1, CS.CEF3 and LD.BER2 (Figure 5A and 5B). Among the visceral tissues, while the cardiac tissue shows the highest TNF-α levels, the cerebral tissue exhibits the lowest TNF-α levels (Figure 5C-5F). Upon treatment by antibiotics, the TNF-α levels are either elevated or declined, implying a complicated interaction among the gut microbiota communities, antibiotics and immune responses. Accordingly, JAK2 and STAT3 that convey an inflammatory signal and initiate cytokine transcription are generally upregulated in the hepatic tissues of CS and LD mice with or without antibiotic supplementation, thereby suggesting an enhanced pro-inflammatory response (Figure 5G and Table 2).

**Figure 5.**
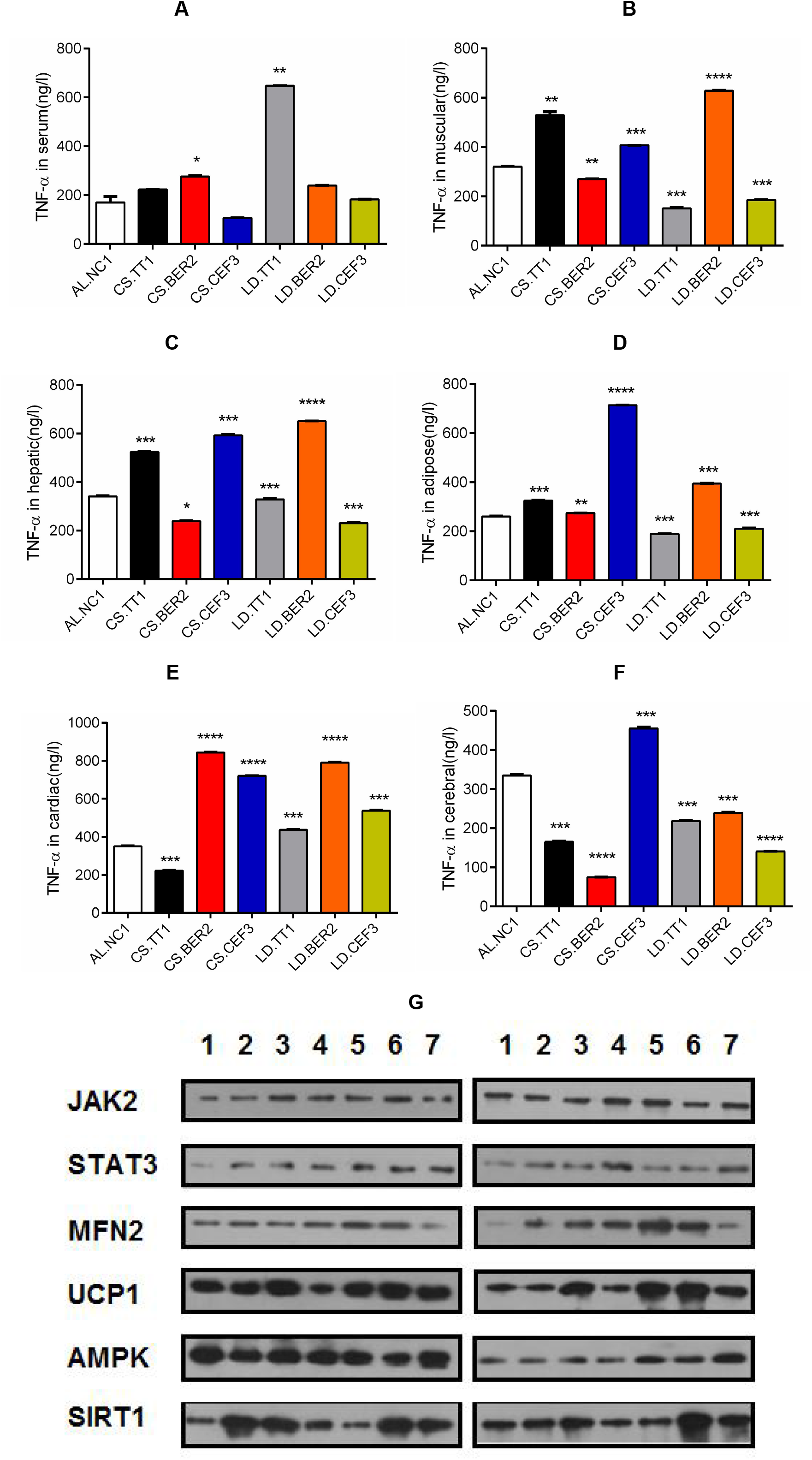
The profiling of pro-inflammatory responses and mitochondrial biogenesis in AL, CS, LD, BER, and CEF mice. (A) TNF-α levels in the serum. (B) TNF-α levels in the muscular tissue. (C) TNF-α levels in the hepatic tissue. (D) TNF-α levels in the adipose tissue. (E) TNF-α levels in the cardiac tissue. (F) TNF-α levels in the cerebral tissue. (G) The hepatic levels (left panel) and muscular levels (right panel) of some critical proteins responsible for mitochondrial biogenesis and mitochondrial biomarker proteins in all tested mice, in which No.1-7 represent AL.NC1, CS.TT1, CS.BER2, CS.CEF3, LD.TT1, LD.BER2, and LD.CEF3, respectively. MFN2: mitofusin 2. UCP1: uncoupling protein 1. AMPK: adenosine 5’-monophosphate-activated protein kinase. JAK2: Janus kinase 2. STAT3: signal transducer and activator of transcription 1 protein. SIRT1: silent mating type information regulation 2 homolog (sirtuin) type 1. GAPDH: glyceraldehyde-3-phosphate dehydrogenase.

**Table 2.** The relative gray scale values of target proteins in the hepatic or muscular tissue of AL, CS, LD, BER, and CEF mice.

|  | AL.NC1 | CS.TT1 | CS.BER2 | CS.CEF3 | LD.TT1 | LD.BER2 | LD.CEF3 |
| --- | --- | --- | --- | --- | --- | --- | --- |
| Hepatic |  |  |  |  |  |  |  |
| JAK2/GAPDH | 0.2469 | 0.3171 | 0.3614 | 0.3580 | 0.3293 | 0.4878 | 0.2561 |
| STAT3/GAPDH | 0.1235 | 0.1829 | 0.2169 | 0.2593 | 0.2927 | 0.3171 | 0.3537 |
| AMPK/GAPDH | 1.5802 | 1.1220 | 1.1807 | 1.1728 | 1.0584 | 0.9390 | 1.4756 |
| SIRT1/GAPDH | 0.4198 | 1.0732 | 1.0482 | 0.6173 | 0.3415 | 1.1951 | 0.8293 |
| MFN2/GAPDH | 0.3210 | 0.3902 | 0.3614 | 0.4691 | 0.5854 | 0.5976 | 0.1829 |
| UCP1/GAPDH | 1.1111 | 1.1098 | 1.4458 | 0.6173 | 1.1585 | 1.5122 | 1.2195 |
| Muscular |  |  |  |  |  |  |  |
| JAK2/GAPDH | 1.0000 | 0.7500 | 0.7179 | 0.9744 | 0.9487 | 0.7750 | 0.9744 |
| STAT3/GAPDH | 0.2564 | 0.3750 | 0.4103 | 0.5385 | 0.3590 | 0.3750 | 0.4615 |
| AMPK/GAPDH | 0.7179 | 0.6750 | 0.7692 | 0.6923 | 1.0513 | 0.9500 | 1.4872 |
| SIRT1/GAPDH | 1.1282 | 1.2250 | 1.3333 | 0.7692 | 0.7436 | 2.4500 | 2.0513 |
| MFN2/GAPDH | 0.2564 | 0.4500 | 1.0256 | 1.0513 | 1.4103 | 1.2750 | 0.3846 |
| UCP1/GAPDH | 0.9744 | 0.7500 | 2.0513 | 0.7692 | 2.3077 | 2.3500 | 1.4615 |

Note: The ratios were calculated from the gray scale value of a target protein dividing that of GAPDH.

To establish an association of pro-inflammation with mitochondrial biogenesis, we quantified some critical proteins responsible for mitochondrial biogenesis and mitochondrial marker proteins in all tested mice. It was clarified that AMPK and SIRT1 transducing an energy deprival signal are globally upregulated in the hepatic and muscular tissues of CS and LD mice with or without antibiotic supplementation although the hepatic tissues exhibit the higher levels than the muscular tissues (Figure 5G and Table 2). Accordingly, MFN2 as a mitochondrial biomarker indicating mitochondrial turnover and UCP1 as an oxidative phosphorylation uncoupling indicator reflecting a conversion from energy deposition to energy expenditure are typically upregulated in the hepatic and muscular tissues of CS and LD mice with or without antibiotic supplementation although give rise to the higher levels in the muscular tissue than those in the hepatic tissue (Figure 5G and Table 2). These results indicated a pro-inflammatory response would induce mitochondrial biogenesis to compensate a functional deficiency of the damaged mitochondria.

### Disordered And Dysfunctional Mitochondria Are Correlated With Augmented Glycolytic product Accumulation And Attenuated Energetic Compound Production

Due to an inflammation-induced hypoxic niche with insufficient oxygen supply, the nascent mitochondria might be dysfunctional, i.e., electron transport along the respiratory chain is interrupted. Under such a condition, mitophagy might be initiated and dysfunctional mitochondria depleted, eventually leading to a net decrease in mitochondrial copy numbers. From the electron microphotographs, more mitochondria were observed in the adipose tissues of AL.NC1 (Figure 6A1) than those in the adipose tissue of other mice (Figure 6A2-6A7). Similarly, more mitochondria were also seen in the muscular tissues of AL.NC1 (Figure 6B1) than those in other mice (Figure 6B2-6B7).

**Figure 6.**
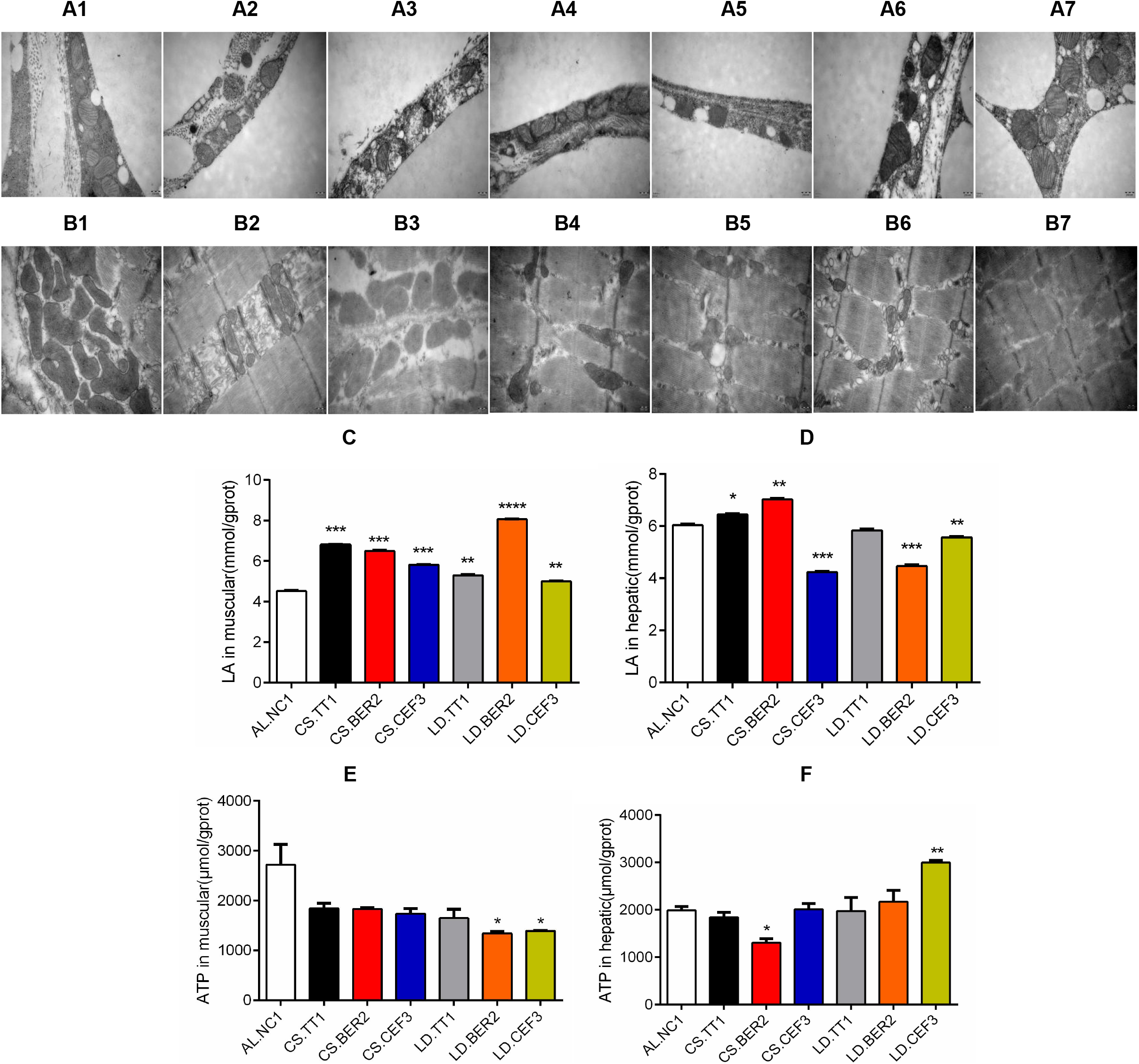
Structure and function of mitochondria in AL, CS, LD, BER, and CEF mice. (A) Mitochondria in the adipose tissues (× 50,000). (B) Mitochondria in the muscular tissues (× 50,000). (C) LA levels in the muscular tissues. (D) LA levels in the hepatic tissues. (E) ATP levels in the muscular tissues. (F) ATP levels in the hepatic tissues. A1-A7 and B1-B7 represent AL.NC1, CS.TT1, CS.BER2, CS.CEF3, LD.TT1, LD.BER2, and LD.CEF3. The bar shows 200 nm in the electron microphotographs.

Accordingly, it was predicted that insufficient or dysfunctional mitochondria should compromise the anaerobic glucose degradation, leading to an elevated lactic acid (LA) level and a declined adenosine triphosphate (ATP) level. As depicted in Figure 6C, the muscular LA levels are globally elevated in all tested mice, but the hepatic LA levels are almost equal to or mildly higher than AL.NC1 (Figure 6D), reflecting a differential efficiency of the functional recovery of mitochondria. Accordingly, a higher muscular ATP level was detected in AL.NC1 compared to other treated mice (Figure 6E), whereas the lowest hepatic ATP level was determined in CS.BER2 among all treatment groups of mice (Figure 6F). These results unraveled a clear correlation of mitochondrial dysfunction with high-level LA and low-level ATP, in which the highest LA and lowest ATP levels in the hepatic tissues of CS.BER2 are coincided to its disturbed gut microbiota status and aggravated pathogenic extent.

### CS Upregulates Atherosclerosis-Related Genes And BER Enhances CS-upregulated Atherogenesis-Related Gene Expression

To monitor the changes from atherogenesis to atherosclerosis upon CS/LD and antibiotic treatments, we quantified the expression levels of 84 atherosclerosis-related genes in AL, CS, LD, BER, and CEF mice (Additional file 2). As compared with AL.NC1 and considering two-fold changes, it was evident that eight genes are upregulated and 21 genes downregulated in CS.TT1 (Figure 7A), while 11 genes are upregulated and 29 genes downregulated in LD.TT1 (Figure 7B). It was noted that *Eng, Itga2, Pdgfa* and *Serpinb2* are commonly upregulated, whereas *Ace, Fgb, Fgf2, Ifng, Il1a, Il1b, Itgax, Ldlr, Lpl, Npy, Sod1* and *Tnfaip3* are generally downregulated in CS.TT1 and LD.TT1. Interestingly, those downregulated genes encode pro-inflammatory cytokines, for example, *Ifng, Il1a, Il1b* and *Tnfaip3* encode IFN-γ, IL-1α, IL-1β and TNF-α.

**Figure 7.**
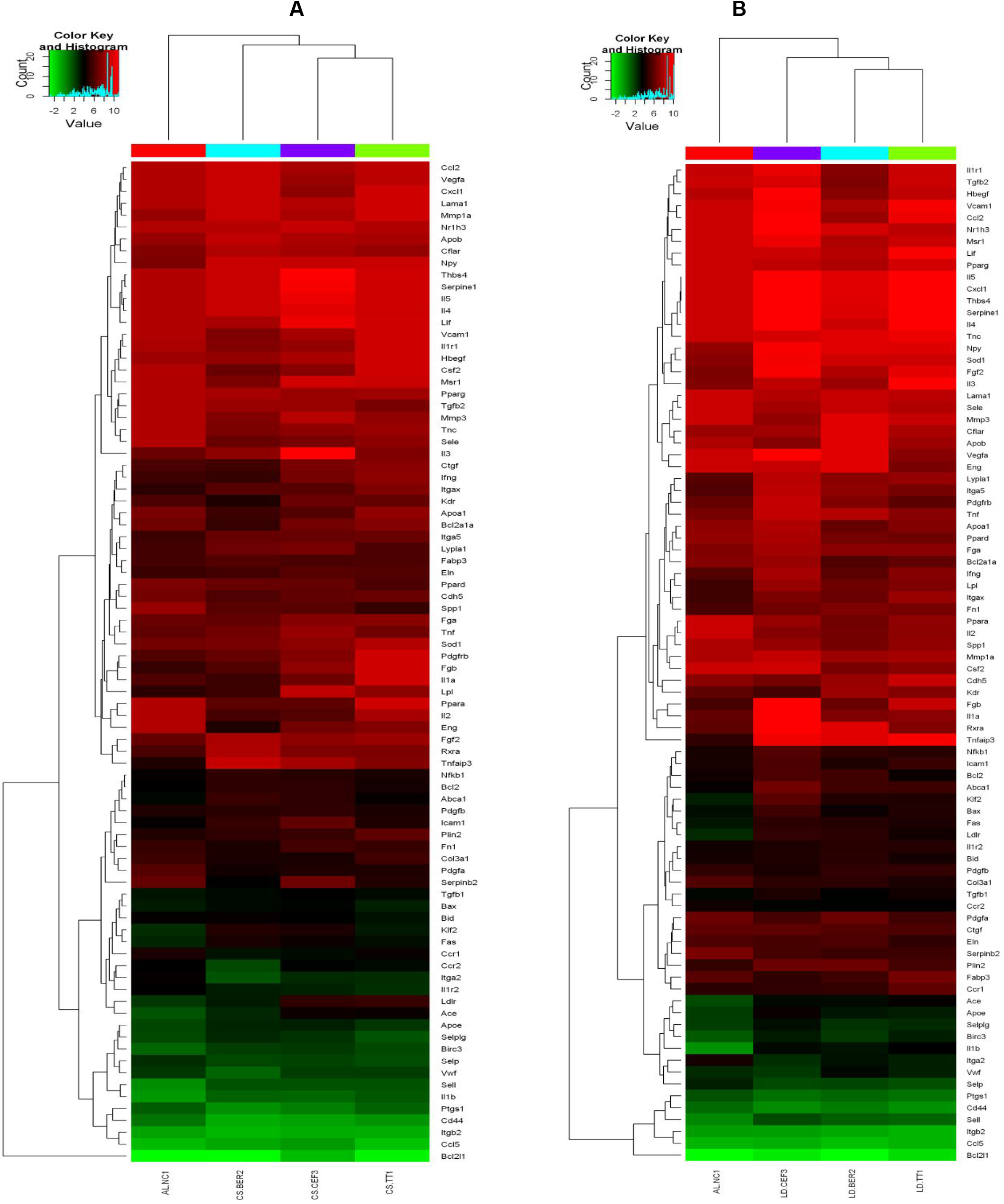
The inflammatory responses and atherosclerosis-related gene expression profiles among AL, CS, LD, BER, and CEF mice. (A) Microarray comparison of 84 atherosclerosis-related genes among the hepatic tissues of AL.NC1, CS.TT1, CS.BER2, and CS.CEF3. (B) Microarray comparison of 84 atherosclerosis-related genes among the hepatic tissues of AL.NC1, LD.TT 1, LD.BER2, and LD.CEF3.

In the CS group, treatment by BER leads to upregulation of 22 genes compared with AL.NC1 and upregulation of 29 genes compared with CS.TT1. In similar, treatment by CEF allows upregulation of 11 genes compared with AL.NC1 and upregulation of 16 genes compared with CS.TT1. In the LD group, 11 and eight genes are upregulated in LD.BER2 and LD.CEF3 compared with AL.NC1, while 19 and 23 genes are upregulated in LD.BER2 and LD.CEF3 compared with LD.TT1 (Table 3). Intriguingly, some pro-inflammatory cytokine, chemokine and corresponding receptor genes are upregulated after treatment by antibiotics. For example, *Ccr1, Csf2, Ifng, Il1a, Il1r1* and *Il2* are upregulated in CS.BER2, and *Ccr1, Csf2, Il1a, Il1r1* and *Il2* are upregulated in CS.CEF3. In similar, *Ccr1, Ifng, Il1r1, Il3, Il4* and *Il5* are upregulated in LD.BER2, and *Ccr1, Il1b, Il1r2, Il2, Il3* and *Il5* are upregulated in LD.CEF3. These results implied antibiotic use enhances pro-inflammatory response, perhaps by inducing more severe gut dysbiosis.

**Table 3.**
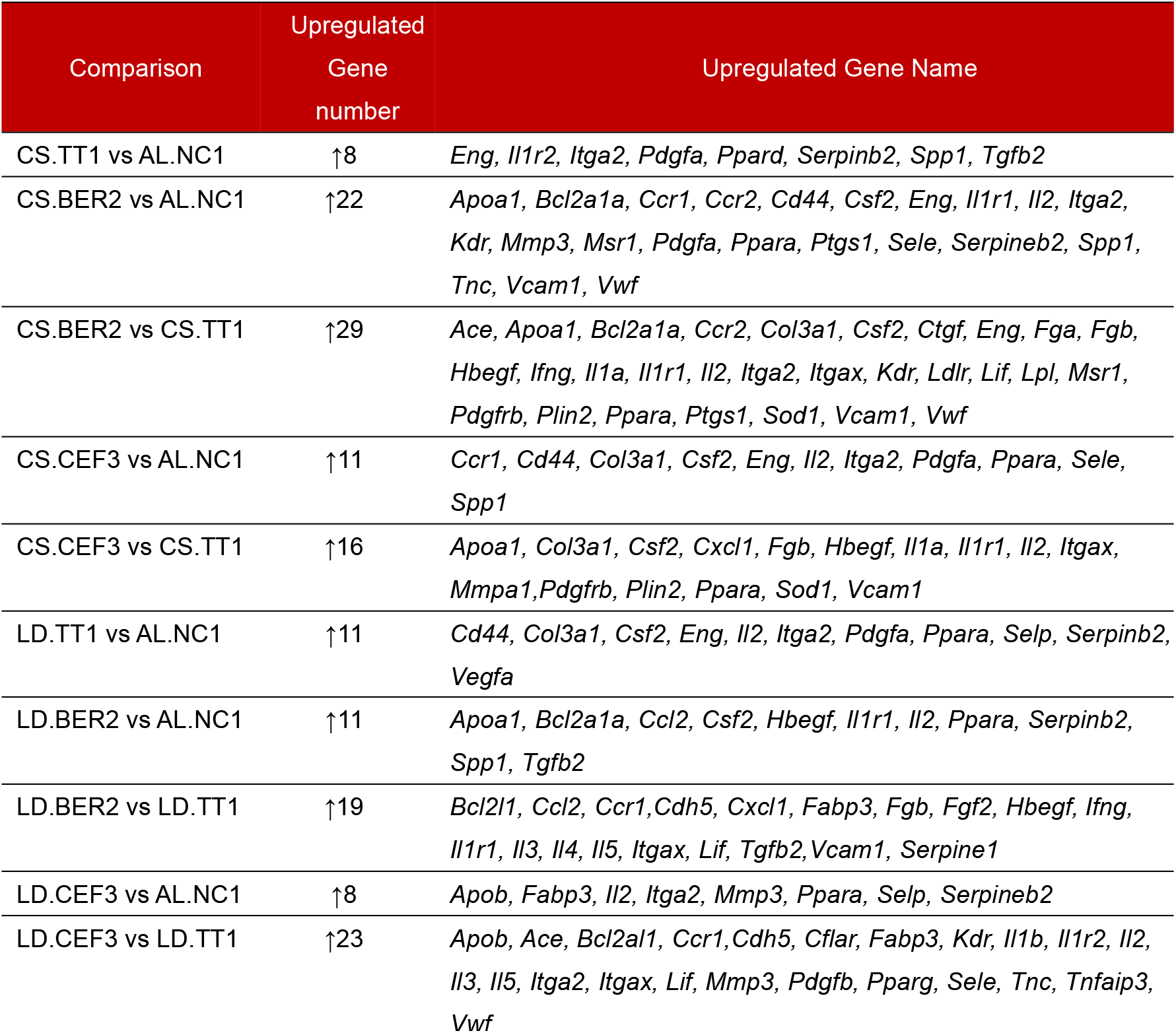
Comparison of atherosclerosis-related genes upregulated for two-fold and above in AL, CS, LD, BER, and CEF mice.

*Vcam1*, upregulated by cytokines in CS.BER2, CS.CEF3 and LD.BER2, encodes a cell adhesion molecule and plays a role in the development of atherosclerosis and rheumatoid arthritis (Wu, 2007). Other frequently upregulated genes also include *Apob, Apoa1, Ppara* and *Eng*. A high level of ApoB was postulated to be the primary driver of plaques that cause vascular disease (atherosclerosis) (Tabas et al., 2007). ApoA1 is often used as a biomarker for prediction of cardiovascular diseases (McQueen et al., 2008). PPAR-α is a transcription factor and a major regulator of lipid metabolism in the liver (Staels et al., 2008). Endoglin is involved in the development of the cardiovascular system and in vascular remodeling (Li et al., 1999).

### CS Induces Steatogenesis And BER Enhances CS-Induced Steatohepatitis-Like Pathogenesis

To reveal a putative association of animal-based diets with hepatic steatogenesis, we fed mice with CS or LD and analyzed the characteristic fatty liver manifestation. It was noticed that the hepatic tissue of AL.NC1 (Figure 8A) exhibits less and small sized oil droplets than that of a CS.TT1 (Figure 8B). Intriguingly, it was also clarified that the more and larger sized hepatic oil droplets emerge in CS.BER2 (Figure 8C), whereas the less and smaller sized hepatic oil droplets occur in CS.CEF3 (Figure 8D). In the LD group, the numbers and volumes of oil droplets are only slightly increased in LD.TT1 (Figure 8E), but dramatically decreased in LD.BER2 (Figure 8F) or LD.CEF3 (Figure 8G). These results indicated BER use for one month can aggravate CS-induced hepatic steatogenesis, but CEF use for the same time only results in less advanced or even attenuated steatogenesis.

**Figure 8.**
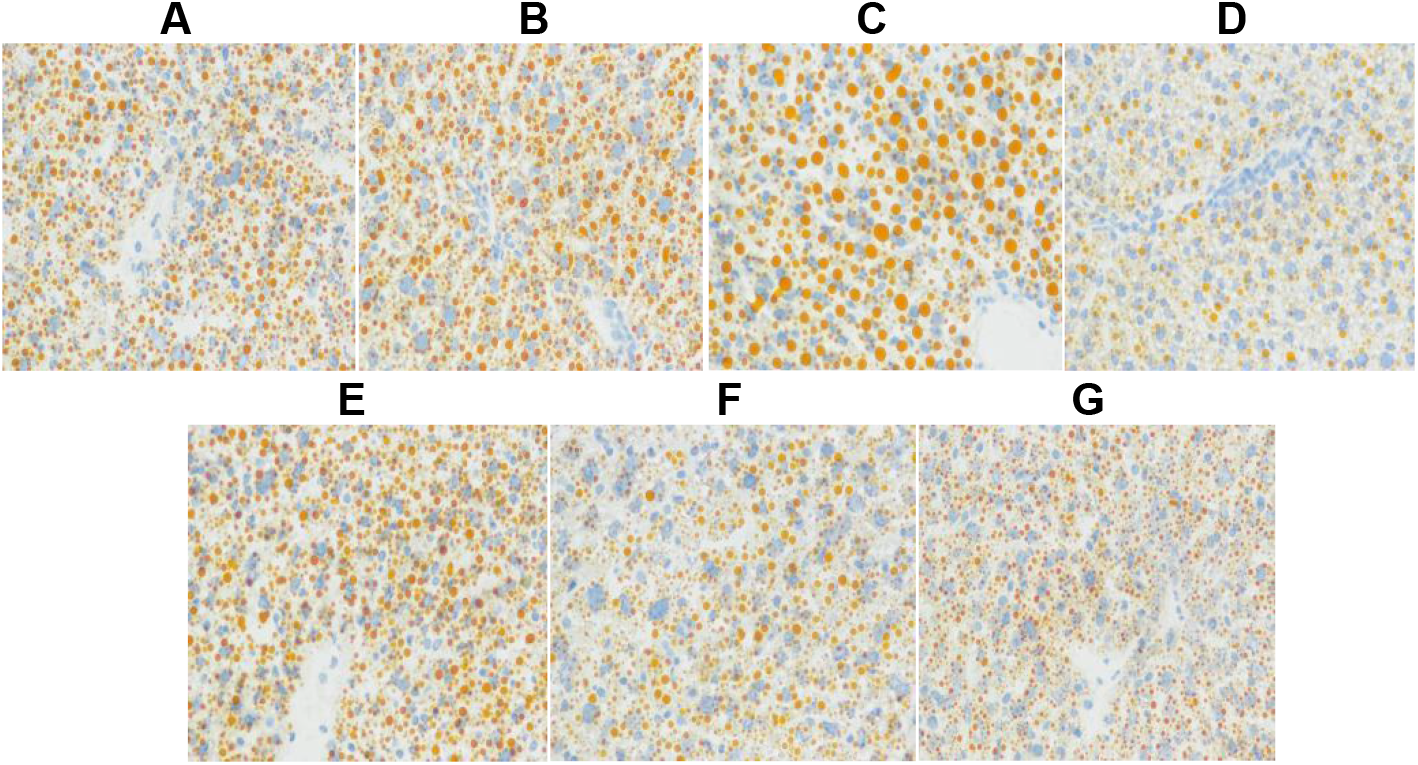
Hepatic steatogenesis and pro-/anti-inflammatory responses in the hepatic and adipose tissues of AL, CS, LD, BER, and CEF mice. (A)-(G) Oil red O staining in the hepatic tissue of AL.NC1 (A), CS.TT1 (B), CS.BER2 (C), CS.CEF3 (D), LD.TT1 (E), LD.BER2 (F), and LD.CEF3 (G), respectively (× 400).

Furthermore, we evaluated the potential risk of CS and CS+BER on NAFLD using 84 genes-comprising mouse fatty liver PCR microarray (Additional file 3). As compared with AL.NC1, while six NAFLD-related genes are upregulated in CS.TT1 with more than two-fold increases of mRNA levels, 12 NAFLD-related genes are upregulated in CS.BER2 with more than two-fold increases of mRNA levels (Table 4). For example, *Serpine 1* mRNA involving in the adipokine signaling pathway is elevated for 15.06 folds in CS.TT1, and 14.83 folds in CS.BER2; *Gsk* mRNA involving in non-insulin dependent diabetes mellitus (NIDDM) is elevated for 3.16 folds in CS.TT1, and 15.67 folds in CS.BER2. Additionally, some upregulated genes in CS.BER2 are downregulated in CS.TT1. For example, *Pdk4, Ldlr*, and *Ppard* are downregulated in CS.TT1, but upregulated in CS.BER2. These results verified CS raises a risk of NAFLD, whereas BER further deteriorates such a severe condition.

**Table 4.**
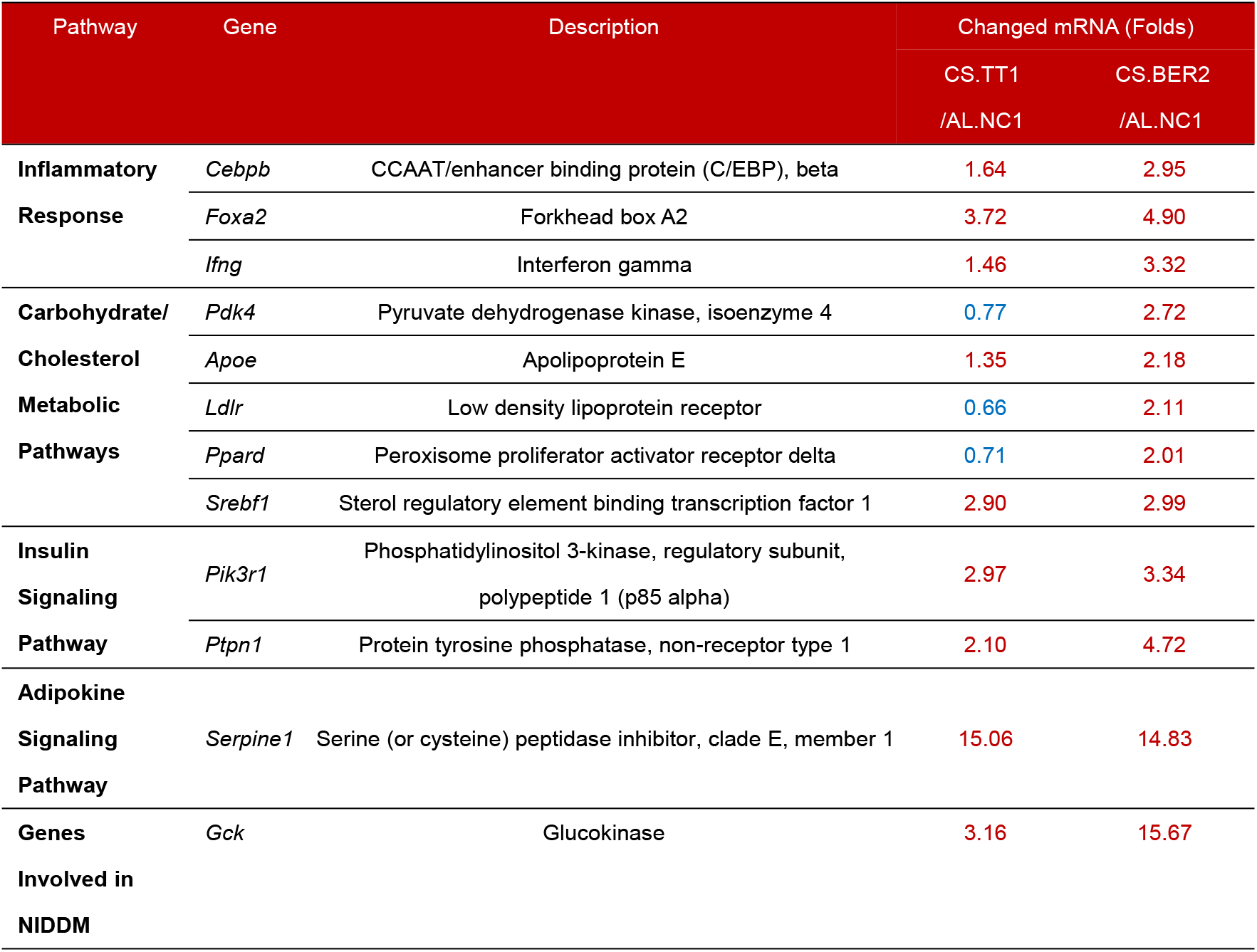
The hepatic expression levels of NAFLD-related genes with up/downregulation for more or less than two folds in AL.NC1, CS.TT1 and CS.BER2 mice

### CS Elevates Dementia Biomarker Levels And BER boosts CS-Induced Dementia Presentation

To evaluate the effects of CS/LD feeding and BER/CEF use on amyloidosis, we determined the level of amyloid β peptide (Aβ), a hallmark of Alzheimer’s disease, in the cerebral tissues of all groups of mice. As results, the cerebral level of Aβ is elevated in CS.TT1, and further elevated to a much higher level in CS.BER2 (Figure 9A). Accordingly, the cerebral level of tyrosine hydroxylase (Th), a rate-limiting enzyme for biosynthesis of the neurotransmitter dopamine, is declined in CS.TT1, and further declined to a much lower level in CS.BER2 (Figure 9B). These results indicated CS increases Aβ and decreases TH, and BER confers a much higher Aβ level and a much lower TH level, thereby implying BER use is a more risk factor than CS feeding in triggering dementia-like pathogenesis with amyloidosis and cognitive deficits.

**Figure 9.**
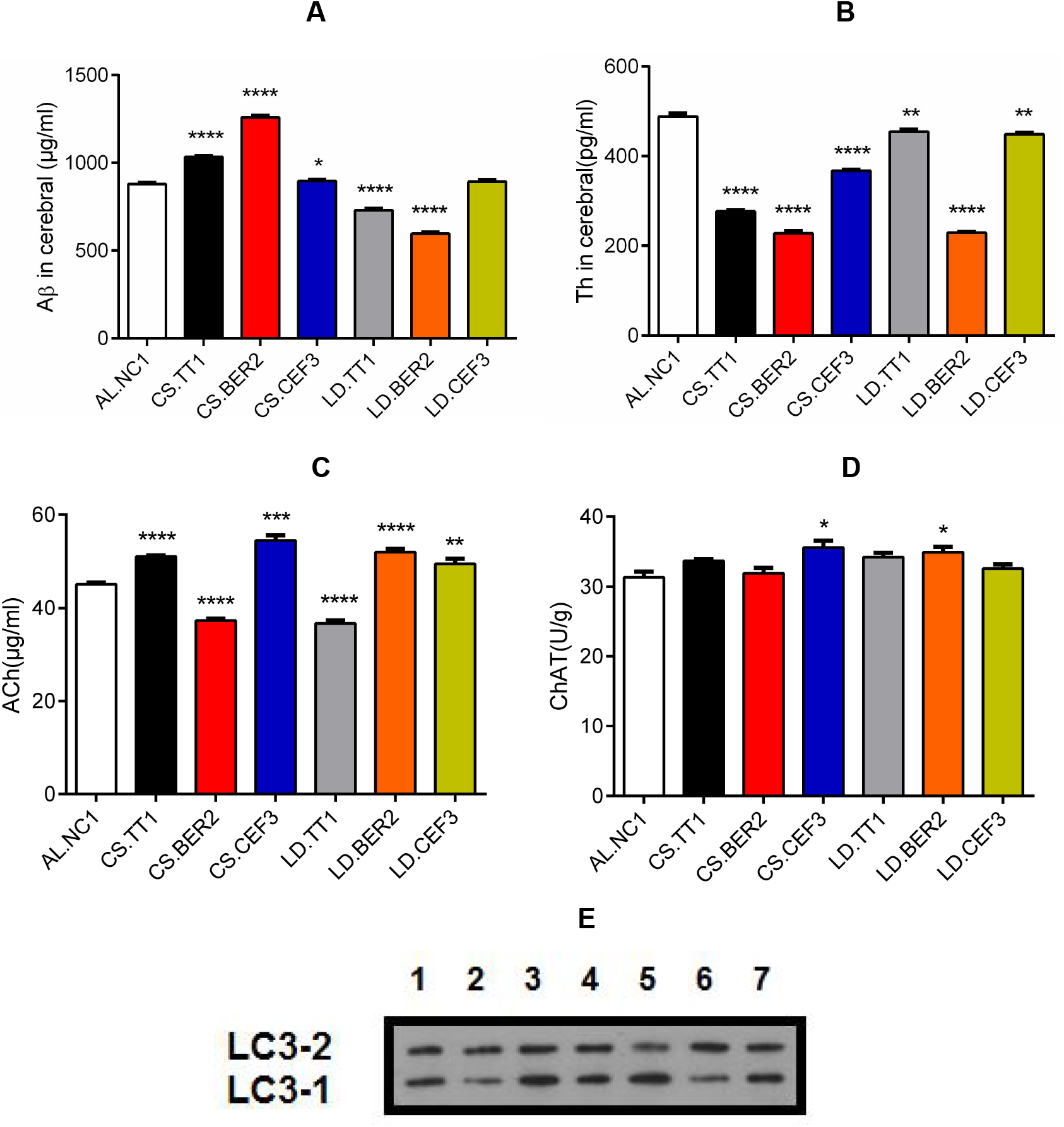
The inflammatory responses and dementia-related gene expression profiles in AL, CS, LD, BER, and CEF mice. (A) Aβ levels in the cerebral tissue. (B) Th levels in the cerebral tissue. (C) Ach levels in the cerebral tissue. (D) ChAT levels in the cerebral tissue. (E) LC3-2/-1 levels in the cerebral tissue, in which No.1-7 represent AL.NC1, CS.TT1, CS.BER2, CS.CEF3, LD.TT1, LD.BER2, and LD.CEF3, respectively.

It was observed that the Aβ level in CS.CEF3 is lower than that in CS.BER2, whereas the Th level in CS.CEF3 is higher than that in CS.BER2, implying an augmented brain pathological alteration in CS.BER2 than in CS.CEF3 (Figure 9A and Figure 9B). On the other hand, elevation of the Aβ level and decline of the Th level were not significantly noted in LD.TT1, LD.BER2 and LD-CEF3 (Figure 9A and Figure 9B), implying no remarkable alterations occurring in the brain structure and function after treatment by LD and/or BER/CEF. Furthermore, CS.BER2 show a low amount of the neurotransmitter acetylcholine (Ach), whereas CS.CEF3 show a high amount of Ach (Figure 9C). Interestingly, LD.BER2 and LD.CEF3 exhibit the higher amounts of Ach than LD.TT1. On the other hand, it was determined that activities of choline acetyltransferase (ChAT) responsible for Ach biosynthesis are almost unchanged among all tested mice (Figure 9D).

To explore the autophagic outcome upon treatments, we quantified microtubule-associated proteins 1A/1B light chain 3B (LC3), the most widely used marker of autophygosomes, and compared the relative level of LC3-1, an unmodified form of LC3, and LC3-2, a lipid modified form of LC3 (Table 4). Consequently, CS.TT1 (lane 2 in Figure 9E), CS.CEF3 (lane 4 in Figure 9E), LD.BER2 (lane 6 in Figure 9E), and LD.CEF3 (lane 7 in Figure 9E) exhibit less LC3-1 and more LC3-2, whereas CS.BER2 (lane 3 in Figure 9E) and LD.TT1 (lane 5 in Figure 9E) show more LC3-1 and less LC3-2. These results suggested a higher autophagic activity (high-rate conversion from LC3-1 to LC3-2) in CS.TT1, CS.CEF3, LD.BER2 and LD.CEF, but a lower autophagic activity (low-rate conversion from LC3-1 to LC3-2) in CS.BER2 and LD.TT1, hence deciphering CS.BER2 possesses a high cerebral Aβ level because it induces a low autophagic activity, and LD.TT 1 possesses a low cerebralAβ level so it also induces a low autophagic activity.

**Table 4.** The relative gray scale values of LC3-1/2 in the hepatic or muscular tissues of AL, CS, LD, BER, and CEF mice.

|  | AL.NC1 | CS.TT1 | CS.BER2 | CS.CEF3 | LD.TT1 | LD.BER2 | LD.CEF3 |
| --- | --- | --- | --- | --- | --- | --- | --- |
| LC3-1/GAPDH (1) | 0.48 | 0.22 | 0.66 | 0.39 | 0.68 | 0.26 | 0.43 |
| LC3-2/GAPDH (2) | 0.42 | 0.35 | 0.50 | 0.47 | 0.34 | 0.64 | 0.49 |
| 1/2 | 1.14 | 0.63 | 1.30 | 0.83 | 2.00 | 0.41 | 0.88 |

Note: The ratios were calculated from the gray scale value of LC3-1/2 dividing that of GAPDH.

## DISCUSSION

A putative association of gut opportunistic infection by commensal bacteria with neurodegenerative diseases was proposed (Li et al., 2016). The clinical manifestation of Parkinson’s disease was shown to positively correlate with the abundance of *Prevotellaceae*, but negatively correlate with the abundance of *Enterobacteriaceae* (Scheperjans et al., 2015). It was demonstrated that oral administration of specific microbial metabolites to germ-free mice promotes neuroinflammation and motor neurodegenerative symptoms, whereas colonization of α-synuclein-overexpressing mice with microbiota from Parkinson’s disease-affected patients enhances physical impairments compared to microbiota transplants from healthy human donors (Sampson et al., 2016). It was revealed that LPS and K99 fimbriae protein (pilin) co-exist in the amyloid plaques of dementia brain tissues, suggesting a cerebral entry of bacterial components (Zhan, et al., 2016). Our study linked the dietary sulfate CS to the mucus-destroying bacteria SSB to the dementia hallmarks, including the high-level Aβ and low-level Th and Ach in the brain. Additionally, we have previously found LPS-containing complete Freund’s adjuvant can trigger brain pathological alterations by increasing Aβ concentrations and decreasing Th activities (He et al., 2017).

A pathogenic mechanism underlying NAFLD or NASH (Brandl & Schnable, 2017) remains unknown although the major risk factors are attributed to the obesity, diabetes and metabolic syndromes (Vuppalanchi et al., 2009). A potential tripartite relationship among the gut, diet, and liver diseases was recognized, a pivotal role of the individual gut microbiome on initiating NAFLD pathogenesis addressed, and animal - or plant-based diets accelerate NAFLD development by modulating the gut microbiome also highlighted (Mokhtari et al., 2017). More importantly, it was believed that sustained LPS leakage elicited by CS might not be completely neutralized by anti-LPS, thereby giving rise to a series of deleterious outcomes, such as induction of diabetes mellitus (Gomes et al., 2017). We found in this study that CS not only enhances adipose storage and impede mitochondrial functions in the liver, but also induces the expression of NAFLD/NIDDM-related genes, implying an early hint of development toward NAFLD/NIDDM.

A correlation of high-fat diets with atherosclerosis was debating and controversial. For example, in a study involving rabbits fed the heated soybean oil, the gross initiation of atherosclerosis and marked induction of liver damage were histologically and clinically demonstrated (Greco & Mingrone, 1990), but it was claimed that this outcome is not caused by dietary cholesterol rather than oxysterols or oxidized cholesterols (Kummerow, 2013). It was known that the dietary component that can be metabolized by the gut microbiota exerts a distinct effect on atherosclerosis, in which the dietary fibre and carnitine play a completely opposite role (Jonsson & Backhed, 2017). Carnitine from red meat and phosphatidylcholine from egg yolk was verified to be converted to trimethylamine by the gut microbiota and subsequently oxidized to trimethylamine N-oxide (TMAO) in the liver, eventually leading to atherosclerosis in the murine models (Wang et al.,2011, Koeth et al.,2013). Our study observed the upregulation of atherosclerosis-related genes by CS feeding, accompanying with deficient and dysfunctional mitochondria.

It was worthy noting from this study that oral CS administration can dramatically change the multi-organ LPS and anti-LPS levels, in which an increase of the serum and muscular anti-LPS level and an accompanying decrease of the serum and muscular LPS level might urge some authors to draw a conclusion that CS exerts an anti-inflammatory effect and is apt to serve as a therapeutic agent for inflammatory diseases (Vergés et al., 2004; Clegg et al., 2006; Reginster et al., 2007). On the other hand, given CS really possesses a therapeutic value for amelioration of OA and IBD, it should only occur in the individuals without SSB, but not happen in the patients with SSB because CS enriches SSB, suggesting evaluation of CS in the inflammatory diseases should consider a potential effect of the gut microbiota.

Whether *A. muciniphila* is beneficial or harmful to human health remains another debating issue although our present work has concluded *A. muciniphila* with an abundance of 0.09% in CS.CEF3 or 0.3% in LD.CEF3 unlikely aggravates the steatohepatitis, atherosclerosis and dementia-like pathogenesis. In similar, other authors have shown antibiotics eliminating *Akkermansia* to reinforce the mucus barrier can block the heme-induced differential expression of oncogenes, tumor suppressors, and cell turnover genes, and prevent the heme-dependent cytotoxic micelles to reach the epithelium (Ijssennager et al., 2015). It was also evident that mucus-eroding microbiota including *A. muciniphila and B. caccae* can promote greater epithelial access and lethal colitis by the mucosal pathogen *Citrobacter rodentium* (Desai et al., 2016). However, *A. muciniphila* was proposed as a novel functional microbe with probiotic properties (Gomez-Gallego et al., 2016).

Indeed, mounting evidence is accumulated regarding the therapeutic values of *A. muciniphila* in metabolic diseases. For example, *A. muciniphila* is inversely correlated with obesity and type I diabetes in both mice and humans (Shin et al., 2014; Dao et al., 2016). The oral administration of *A. muciniphila* prevents a high-fat diet-induced obesity by altering adipose tissue metabolism and gut barrier function (Everard et al., 2013). The anti-obesity effect of a polyphenol-rich cranberry is associated with increased *Akkermansia* spp. population in the mouse gut (Anhê et al., 2015). Recently, a favorable effect of *A. muciniphila* on the improvement of glucose tolerance has been revealed upon maintaining a high IFNγ level for controlling the multiple and primarily intracellular pathogen infections (Greer et al., 2016). Similarly, a positive correlation between A. *muciniphila* and gut health has been partially explained by its immune modulatory effects and immune tolerance to this commensal (Ottman et al., 2017).

As our opinion, there might be a threshold effect impacting the outcome of A. *muciniphila*, in which a minimal abundance of A. *muciniphila* might induce mucin production and promote mucus repair because it is anchored on the mucus and thus able to efficiently activate the mucus immunity to combating the gut opportunistic infection, but a maximal abundance of A. *muciniphila* should allow the excessive mucus opening and large-scale LPS leakage into the blood stream. In our present study, CEF was found to eradicate *Desulfovibrio*, but not to kill *A. muciniphila*, thereby exhibiting a less deleterious impact on the systemic pathogenesis under investigation. In contrast, we found BER use for one month can completely kill *A. muciniphila*, but cannot eradicate *Desulfovibrio*, showing an augmenting role on the systemic inflammatory pathogenesis. These results are well consistent with the popular concept of A. *muciniphila* as a probiotic beneficial to human health (Gomez-Gallego, et al., 2016).

BER as an antibacterial agent has been demonstrated to inhibits the growth of *Staphylococcus aureus* (Stermitz et al., 2000) and *Microcystis aeruginosa* (Zhang et al., 2010) when applied *in vitro* in combination with methoxyhydnocarpin, an inhibitor of multidrug resistance pumps. Some researchers have also explored the possible application of BER against the tractable infection by multi-resistant *Staphylococcus aureus* (MRSA) (Yu et al., 2005). Additionally, BER is under investigation to determine whether it may have applications as a drug in treating diabetes, hyperlipidemia, and cancer. Recently, BER has been proven to improve glucogenesis and lipid metabolism in NAFLD (Zhao et al., 2017), perhaps mainly acting on the visceral organs in addition to the gut microbiota *per se*.

Overall, it is likely impossible to eradicate SSB/SRB from the gut by the monotherapy with a single antibiotic even a long-term use although CEF incidentally ameliorates CS/LD-driven systemic inflammatory disorders by nursing the beneficial probiotic bacterium. In such a context, a homeostatic gut symbions is a pivotal determinant for human gut health. Some studies have unraveled the diet, exercise and antiglycemic drugs may alter the development towards NFALD/NASH (Adams & Angulo, 2006; Veena et al., 2014), but no pharmacological treatment regimens have been approved until 2015 (Ratziu et al., 2015). It is generally believed that a reasonable replacement of red meat by fruits and vegetables can impede the progression of atherosclerosis and decrease the risk of cardiovascular diseases (Wang et al., 2014). Our present study also addressed a fact that the meat diet rich in CS can evoke the systemic inflammatory pathogenesis by enriching mucus-destroying bacteria, strongly advising to restrict daily meat intake as early and frequently as possible.

## EXPERIMENTAL PROCEDURES

### Animals, Diets And Antibiotic Uses

The female Kunming mice (four-week old, 15-20 g) that belong to an out-bred population from SWISS mice were provided by The Experimental Animal Centre of Guangzhou University of Chinese Medicine in China (Certificate No. 44005800001448). All mice were housed on a 12-h light and 12-h dark cycle at 25°C with *ad libitum* chow and free water drinking. After one-week quarantine, mice were randomly divided into four groups: a control group of mice fed with *ad libitum* chow (AL.NC1), a model group of mice intragastrically fed with 250 μl CS (100 mg/ml, FocusChem, Shandong, China) daily for one month in addition to *Ad libitum* (CS.TT1), a model group of mice intragastrically fed with 200 μl warm lard daily for one month in addition to *Ad libitum* (LD.TT1), a treatment group of CS mice administered BER via drinking water containing 0.1 g/ml of berberine hydrochloride (CS.BER2) or containing 100 mg/ml of cefotaxime sodium (CS.CEF3) for one month, and a treated group of LD mice administered BER via drinking water containing 100 mg/ml of berberine hydrochloride (LD.BER2) or containing 100 mg/ml of cefotaxime sodium (LD.CEF3) for one month. Three to six mice were included within each group. All experiments were approved by The Animal Care Welfare Committee of Guangzhou University of Chinese Medicine (No. SPF-2015009). The experimental protocols complies with the requirements of animal ethics issued in the Guide for the Care and Use of Laboratory Animals of the NIH, USA.

### Gut Microbiota Metagenomic Analysis And Gene Expression Microarray

The gut microbiota in a mouse fecal sample was identified by the high-throughput 16S VX sequencing-based classification procedure. The sequencing (sample preparation, DNA extraction and detection, amplicon purification, library construction and online sequencing) and data analysis (paired end-reads assembly and quality control, operational taxonomic units cluster and species annotation, alpha diversity and beta diversity) were conducted by Novogene (Beijing, China). The RT^2^ Profiler™ PCR Array Mouse Atherosclerosis (PAMM-038Z) and the RT^2^ Profiler™ PCR Array Mouse Fatty Liver (PAMM-157Z) were purchased from SABioscience Qiagen (Hilden, Germany). The experiment was performed by Kangchen Biotechnology (Shanghai, China).

### Western Blotting (WB) And Enzyme-Linked Immunosorbent Assay (ELISA)

The reference protein GAPDH and all antigen proteins were immunoquantified by WB according to manufacture’s manuals. The gray scale values were analyzed by the IMAGEJ software. Antibodies against JAK2, STAT3, AMPKα1, SIRT1, MFN2, UCP1 were purchased from Abcam (Cambridge, UK). ELISA kits for TNF-α, LPS, anti-LPS, and TH were purchased from Chenglin Biotech (Beijing, China).

### Histochemical Analysis

Cut fresh frozen tissue sections at 5-10 μm thick and mount on slides. Air dry slides for 30-60 min at room temperature and then fix in ice cold 10% formalin for 5-10 min. Air dry again for another 30-60 min or rinse immediately in 3 changes of distilled water. Let slides air dry for a few min. Place in absolute propylene glycol for 2-5 min to avoid carrying water into oil red O. Stain in pre-warmed 0.5% oil red O solution (0.5 g oil red O and 100 ml propylene glycol) for 8-10 min in 60 °C oven. Differentiate in 85% propylene glycol solution for 2-5 min. Rinse in 2 changes of distilled water. Stain in haematoxylin for 30 sec. Wash thoroughly in running tap water for 3 min. Place slides in distilled water. Mount with glycerin jelly.

### Immunohistochemical Analysis

The primary antibody against SLPI was purchased from Novus Biologicals (Littleton, USA), and the primary antibody against MUC1 was provided by CapitalBio (Beijing, China). Formalin-fixation, paraffin-embedment and deparaffinization were the same as HE staining procedure described above. Sections were incubated at room temperature with 3% H_2_O_2_ to block endogenous peroxidase, and then repaired in boiling citric acid. After washing in phosphate-buffered solution, sections were blocked by 2% bovine serum albumin and incubated with 1:100 diluted primary antibodies at 37°C for 1 h. After washing again, sections were incubated with biotinylated secondary antibodies at 37°C for 20 min. After washing again, sections were incubated with diaminobenzidine for 1-5 min. After rinsing with tap water, sections were counter stained by haematoxylin. After completion of dehydration, clearance and mounting, photos were taken under the microscope (OLYMUPUS BX-51).

### Electronic Microscopy and Laser Con-focal Microscopy

After treatment, cells were harvested and fixed in 2.5% glutaraldehyde in 0.1 M phosphate buffer for 3 h at 4 °C, followed by post-fixation in 1% osmium tetroxide for 1 h. Samples were dehydrated in a graded series of ethanol baths, and infiltrated and embedded in Spurr’s low-viscosity medium. Ultra-thin sections of 60 nM were cut in a Leica microtome, double-stained with uranyl acetate and lead acetate, and examined in a Hitachi 7700 transmission electron microscope at an accelerating voltage of 60 kV.

The fresh samples of mouse skeletal muscles were fixed in paraformadehyde for 24 h. After repeatedly rinsed, the fixed tissues were dehydrated by gradient ethanol. For embedding and sectioning, the tissue slices were pasted on the slides and coated at 50 °C. After they were de-waxed by a transparent reagent and rinsed, the slides were incubated with antibodies and stained by 4’,6-diamidino-2-phenylindole (DAPI). After drying, a fluorescence quencher was added and slides were sealed. The fluorescence-labeled second antibodies and the first antibody against MUC1 were purchased from Biosynthesis Biotechnology (Beijing, China). Immunoblotting was carried out based on the manufacturer’s instructions.

### Spectrophotometry

The reagent kits for measurement of ATP and LA were purchased from Jiancheng Biotech (Nanjing, China). All determination procedures were according to manufacturers’ instructions.

### Statistical Analysis

The software SPSS 22.0 was employed to analyze data, and the software GraphPad Prism 5.0 was employed to plot graphs. The Independent Simple Test was used to compare all groups, but the Kruskal-Wallis Test followed by Nemenyi test was used when the data distribution is skewed. The significance level (*p* value) was set at <0.05 (*), <0.01 (**), <0.001 (***) and <0.0001 (****).

## SUPPLEMENTARY INFORMATION

The datasets supporting the conclusions of this article are included in the supplementary information (**Additional file 1** includes gut microbiota metagenomic data, **Additional file 2** includes atherosclerosis-related gene microarray data, and **Additional file 3** includes fatty liver-related gene microarray data).

## AUTHOR CONTRIBUTIONS

QPZ, QW and QX designed the study. TL, YPC, and LLT carried out the experiments. XAH performed bioinformatics analysis. SQH, CQL, QW and QX participated in the interpretation of results. QPZ wrote the manuscript with input from other authors. All authors read and approved the final manuscript.

## ACKNOWLEDGMENTS

This work was supported by the National Natural Science Foundation of China (No. 81273620 to Qing-Ping Zeng, No. 81673861 to Chang-Qing Li, and No. 81273817 and No. 81473740 to Qi Wang), and Guangdong Science and Technology Plan Project (No. 20150404042 to Qin Xu). We thank our colleagues in the Tropical Medicine Institute and Clinical Pharmacology Institute, Guangzhou University of Chinese Medicine, China. We also thank Kangchen Biotechnology Co, Shanghai, China for performance of RT-PCR microarray experiments, and Novogene, Beijing, China for conduction of gut microbiota metagenomic analysis.

